# Integration of vascular progenitors into functional blood vessels represents a novel mechanism of vascular growth

**DOI:** 10.1101/2021.11.09.467331

**Authors:** Sanjeeva Metikala, Michael Warkala, Satish Casie Chetty, Brendan Chestnut, Elizabeth Plender, Olivia Nester, Sophie Astrof, Saulius Sumanas

## Abstract

During embryogenesis, the initial vascular network is thought to form by the process of vasculogenesis, or the specification of vascular progenitors *de novo*. After the initial blood circulation has been established, the majority of later-forming vessels are thought to arise by angiogenesis from the already established vasculature. Here we show that new vascular progenitors in zebrafish embryos contribute to functional vasculature even after blood circulation has been established. Based on the expression analysis of early vascular progenitor markers *etv2* and *tal1*, we characterized a novel site of late vasculogenesis (termed secondary vascular field, SVF), located bilaterally along the yolk extension. Using time-lapse imaging of *etv2* reporter lines, we show that SVF cells migrate and incorporate into functional blood vessels and contribute to the formation of the posterior cardinal vein and subintestinal vasculature, suggesting a novel mode of vascular growth. We further demonstrate that SVF cells participate in vascular recovery after chemical ablation of vascular endothelial cells. Inducible inhibition of *etv2* function prevented SVF cell differentiation and resulted in the defective formation of subintestinal vasculature. In addition, we performed single-cell RNA-seq analysis to identify the transcriptional profile of SVF cells, which demonstrated similarities and differences between the transcriptomes of SVF cells and early vascular progenitors. Our results characterize a novel mechanism of how new vascular progenitors incorporate into established vasculature and revise our understanding of basic mechanisms that regulate vascular development.

## Introduction

The vascular network is one of the earliest organ systems to form in a developing vertebrate embryo. It plays a central role in supplying oxygen and nutrients, maintaining tissue homeostasis and is also implicated in numerous diseases. Vascular development is a tightly regulated process and is highly conserved among vertebrates (Beis and Stainier, 2006; Isogai et al., 2001; Lawson and Weinstein, 2002a). The formation of a vascular network occurs in two distinct morphogenetic processes. The *in situ* aggregation of angioblasts, also known as vascular endothelial progenitors into vascular cords that are remodeled into functional blood vessels, is termed vasculogenesis (Patan, 2000; Poole and Coffin, 1989; Risau et al., 1988). The earliest blood vessels which include the dorsal aorta (DA) and posterior cardinal vein (PCV) are known to form by the process of vasculogenesis from angioblasts that originate in the lateral plate mesoderm (LPM) (Risau et al., 1988). Some of the later forming vessels which include pharyngeal arch vessels (Wang et al., 2017) and mammalian organ-specific vasculature (Gouysse et al., 2002; Matsumoto et al., 2001) are also thought to form by vasculogenesis. In the subsequent phase, new vessels grow from pre-existing vessels by sprouting and intussusception, referred to as angiogenesis (Risau et al., 1988; Patan, 2000; Makanya et al., 2009). The current dogma is that vasculogenesis is largely limited to the formation of the initial vascular network during embryogenesis, while the majority of the later vessels form by sprouting from the existing vessels. This view has been recently challenged by the evidence that erythro-myeloid progenitors can contribute to the forming vascular network (Plein et al., 2018). Although additional sources of vascular progenitors in adults have been reported including a vasculogenic zone in the adult human internal thoracic artery vascular wall (Zengin et al., 2006) and circulating endothelial progenitor cells in human peripheral blood derived from bone-marrow (Asahara et al., 1997; Takahashi et al., 1999), their contribution to adult vasculature is still debated (Yoder, 2012). It is currently unknown whether there are additional sources of new vascular progenitors that may contribute to the existing functional vasculature.

While it can be challenging to visualize early vascular development in mammalian embryos, the zebrafish has emerged as an advantageous model system for studying vascular network formation in recent years. Zebrafish embryos possess several features such as optical clarity and high genetic amenability, which, coupled with the availability of fluorescent transgenic lines, enables relatively easy visualization of morphogenetic events in live embryos. Importantly, the molecular mechanisms that regulate vasculogenesis and angiogenesis are highly conserved between zebrafish and mammals (Ellertsdóttir et al., 2010). In zebrafish, vasculogenesis commences as early as 10-11 hours post fertilization (hpf). Angioblasts emerge from the LPM and migrate to the midline where they coalesce to form the DA and PCV by the 22-somite stage (∼20 hpf) (Eriksson and Löfberg, 2000; Herbert et al., 2009; Kohli et al., 2013; Lawson and Weinstein, 2002b; Williams et al., 2010). Subsequently, intersegmental vessels (ISVs) begin to sprout from the DA at ∼22 hpf in the zebrafish trunk shortly before the initiation of blood circulation at 24 hpf, followed by the formation of the dorsal longitudinal anastomosing vessel (DLAV) at 30 hpf (Bahary et al., 2007; Covassin et al., 2006; Hogan and Schulte-Merker, 2017).

Recent studies have demonstrated that several different types of progenitor cells emerge from the PCV, including progenitors of venous ISVs, lymphatic progenitors and organ-specific vascular progenitors, which migrate and coalesce into the subintestinal artery (SIA) and subintestinal vein (SIV) (Hen et al., 2015; Koenig et al., 2016; Nicenboim et al., 2015). It is currently debatable if these different subtypes of endothelial cells are derived from different pools of vascular progenitors which are prespecified during early stages of vasculogenesis, or whether they all share the same type of common progenitor.

We and others have previously identified an ETS transcription factor, *Etv2* / *Etsrp* / *ER71* which functions as an evolutionarily conserved critical regulator of vasculogenesis (Ferdous et al., 2009; Lee et al., 2008; Pham et al., 2007; Sumanas and Lin, 2005). It is one of the earliest markers expressed in vascular endothelial progenitor cells. Both zebrafish and mammalian embryos deficient in Etv2 function fail to form functional vasculature vasculogenesis (Ferdous et al., 2009; Lee et al., 2008; Pham et al., 2007; Sumanas and Lin, 2005). While *etv2* is strongly expressed in vascular progenitors, its expression is downregulated after the establishment of functional vasculature at ∼24 hours post fertilization (hpf) in zebrafish embryos.

In the current study, we characterized a novel site of vasculogenesis, the secondary vascular field (SVF), along the yolk extension in the zebrafish trunk. We showed that SVF-derived vascular endothelial progenitor cells incorporate into the existing vasculature and contribute to the formation of subintestinal vessels. Inducible inhibition of *etv2* function prevented the differentiation of SVF-derived endothelial cells and resulted in the defective formation of subintestinal vasculature. After all other vascular endothelial cells were ablated, SVF cells were sufficient for partial regeneration of the vascular system. Our findings describe a novel mechanism that contributes to vascular growth which will be important for understanding normal and pathological vascular development.

## Results

### New vascular endothelial cells contribute to blood vessels after the initiation of circulation at 24 hpf in zebrafish

To test if new vascular endothelial cells can arise within the trunk region of zebrafish embryos after the initiation of blood circulation at 24 hpf, we utilized a previously generated *TgBAC(etv2:Kaede)* (Kohli et al., 2013) and a newly generated *Tg(fli1a:Kaede)* reporter lines, which express a photo-convertible Kaede protein in vascular endothelial cells driven by the *etv2* and *fli1a* promoters, respectively. We performed a complete photo-conversion of the Kaede protein from green to red at 24 hpf (Figure S1) and imaged the trunk region at 48 and 72 hpf. Vascular endothelial cells (VECs) that were specified before photo-conversion appeared both red and green whereas new VECs that emerged after photo-conversion appeared only green. We observed the majority of newly specified VECs in the PCV, SIV and SIA (Figure 1). A few green only cells were also located in the DA, ISVs and DLAV (Figure 1P,Q). Interestingly, many of the VECs that contributed to the SIV and SIA exhibited only green Kaede expression, suggesting that they originated after the photoconversion (Figure 1D,E,I,J,N,O).

**Figure 1.**
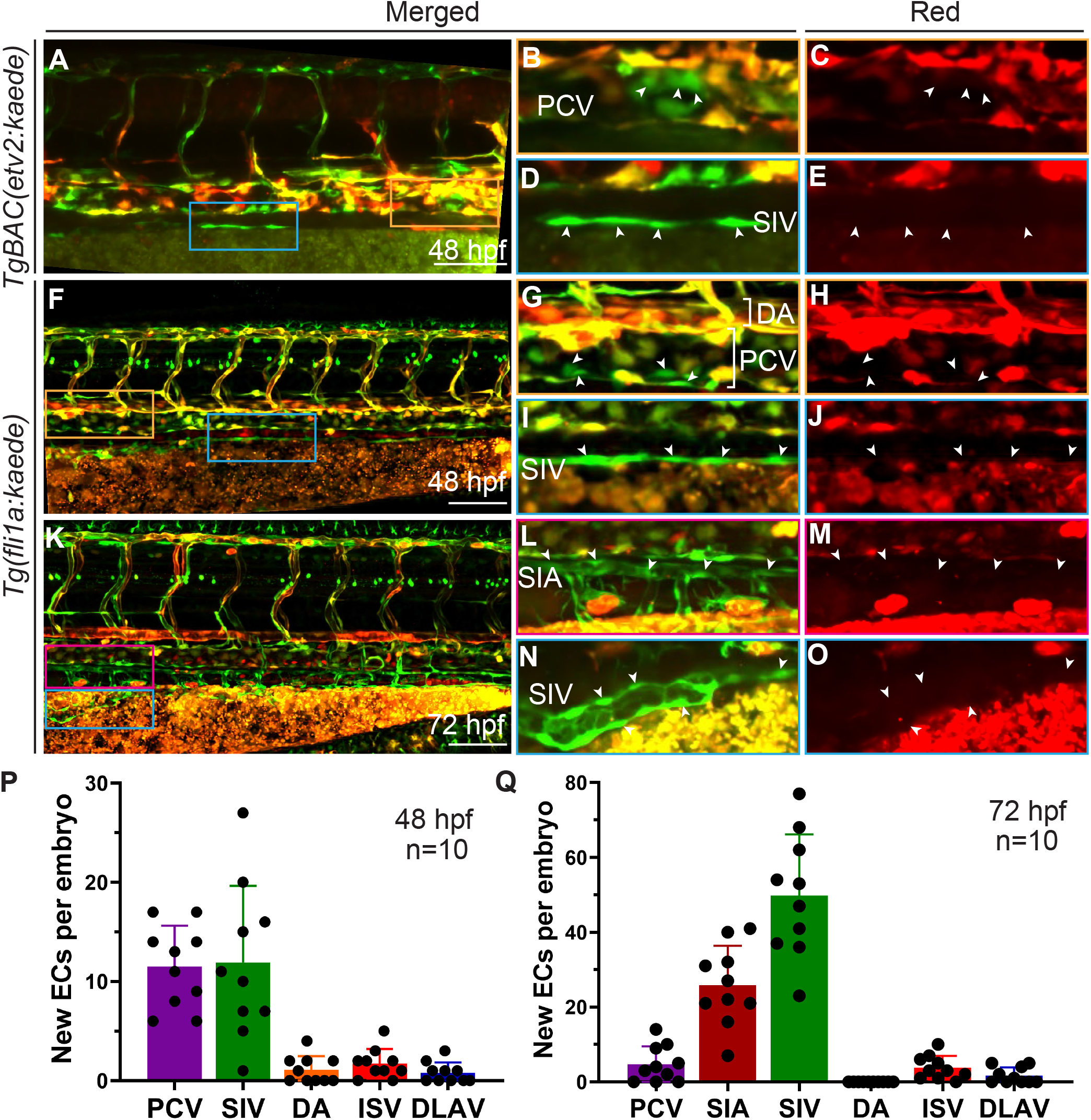
New cells contribute to blood vessels after 24 hpf: (A-O) *TgBAC(etv2:Kaede)* and *Tg(fli1a:kaede)* embryos were photo-converted from green to red at 24 hpf. New vascular endothelial cells were observed in the trunk region at 48 and 72 hpf in PCV (orange insets, B,C,G,H), SIV (blue insets, D,E,I,J) and SIA (magenta inset, L,M). White arrowheads show selected VECs which are green only and do not have red *Kaede* fluorescence. Maximum intensity projections of confocal z-stack images are shown except for magnified insets which show a projection of limited number of z planes. (P,Q) Quantification of new green only VECs using *Tg(fli1a:kaede)* embryos at 48 (n=10) and 72 hpf (n=10) respectively. 10 out of 10 embryos showed both green only and green + red cells. mean ± s.d. Each data point represents the number of green only cells per embryo. Data from 4 independent experiments. PCV, posterior cardinal vein; DA, dorsal aorta; ISV, inter-segmental vessels; SIV, subintestinal vein; SIA, supraintestinal artery. Scale bars: 100 µm.

### Identification of a secondary vascular field (SVF)

To identify these late-forming vascular progenitors, we performed whole mount *in situ* hybridization (WISH) analysis of vascular endothelial and hematopoietic progenitor markers, *etv2* and *scl / tal1* at 24-48 hpf stages. As expected, *etv2* expression was observed in the vasculature and *tal1* expression was observed in blood cells, neural tube and weakly in blood vessels (Gering et al., 1998; Sumanas and Lin, 2005). In addition, bilaterally positioned cells with strong *etv2* and *tal1* expression were seen along the yolk extension outside of the established axial vasculature (Figure 2A,C). These cells were observed from 24 hpf to 48 hpf, and their number reached a maximum between 30 and 36 hpf (Figure 2B,D). Some of these cells appeared closer to the PCV than others, suggesting migratory behavior and the potential to contribute to the vasculature. A double fluorescent *in situ* hybridization (FISH) against both *etv2* and *tal1* showed co-expression of both markers in the same cells (Figure 2E). Although previous studies have reported bilateral *etv2* (Sumanas and Lin, 2005) and *tal1* (Bussmann et al., 2007) expression in similar cells positioned along the yolk extension adjacent to the pronephros, their identity and function have not been investigated further. To confirm the anatomical location of these *etv2+tal1+* cells, we performed fluorescent *in situ* hybridization (FISH) against *etv2*, combined with fluorescence analysis for the pronephric duct marker *Tg(enpep:GFP)* (Seiler and Pack, 2011). Indeed, *etv2+* cells were positioned either medially, laterally or ventrally adjacent to the pronephros (Figure 2F,G and Video S1). Furthermore, we also observed expression of *lmo2* in SVF cells, which similar to *etv2* and *tal1,* is known to label vascular endothelial and hematopoietic precursors (Thompson et al., 1998). In contrast, *fli1a*, a marker of differentiated vascular endothelial cells (Thompson et al., 1998), was absent from SVF cells (Figure S2). We hypothesized that these cells may be late-forming vascular progenitors, and therefore we named this region as the Secondary Vascular Field (SVF).

**Figure 2.**
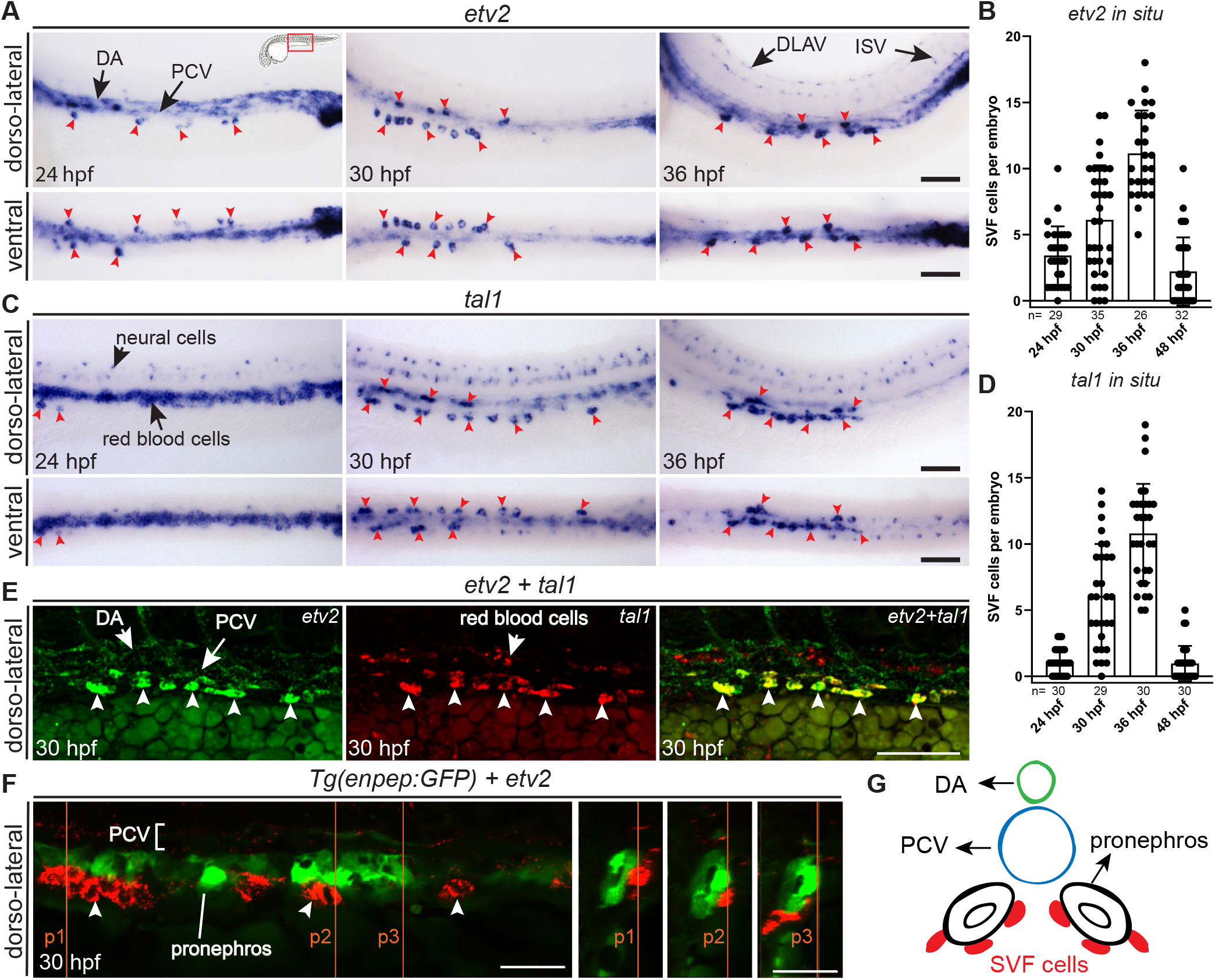
*Etv2* and *tal1* expression in the presumptive secondary vascular field (SVF): (A) Whole-mount *in situ* hybridization (WISH) in wild-type (wt) embryos showing *etv2* expression from 24 to 36 hpf. Cells with strong *etv2* expression were observed in the trunk region, positioned bilaterally along the yolk extension (red arrowheads). Weaker *etv2* expression was also observed in the established vasculature including the DA, PCV, DLAV and ISVs. Dorso-lateral and ventral views of the same embryos are shown. (B) Quantification of SVF cells expressing *etv2* (24 hpf, n=29; 30 hpf, n=35; 36 hpf, n=26 and 48 hpf, n=32) from 24-48 hpf, mean ± s.d. Each data point represents the number of cells in a single embryo. Data from 2 independent experiments. (C) WISH in wt embryos showing *tal1* expression from 24 to 36 hpf. Cells with strong *tal1* expression were observed in the trunk, positioned bilaterally along the yolk extension (red arrowheads). In addition, *tal1* is expressed in red blood cells and a subset of neurons within the spinal cord. Dorsolateral and ventral views of the same embryos are shown. (D) Quantification of SVF cells expressing *tal1* (24 hpf, n=30; 30 hpf, n=29; 36 hpf, n=30 and 48 hpf, n=30) from 24-48 hpf, mean ± s.d. Each data point represents the number of cells in a single embryo. Data from 2 independent experiments. (E) Multiplex HCR *in situ* showing co-expression of *etv2* and *tal1* in SVF cells (white arrowheads). (F) Fluorescent *in situ* hybridization (FISH) against *etv2* (red) combined with the pronephric marker *Tg(enpep:GFP)* fluorescence (green) shows that SVF cells are positioned next to the pronephros in a zebrafish embryo at 30 hpf. Dorso-lateral and transverse confocal sections at selected points (p1, p2 and p3) are shown. (G) Transverse section diagram showing the position of SVF cells with respect to the pronephros and PCV at 30 hpf. PCV, posterior cardinal vein; DA, dorsal aorta; ISV, intersegmental vessels; DLAV, dorsal longitudinal anastomosing vessel. Scale bars: A, B: 100 µm, D: 50 µm, E: 25 µm.

### SVF cells contribute to the PCV and subintestinal vasculature

To visualize SVF cells in live embryos, we utilized the CRISPR/Cas9-mediated non-homologous integration approach (Auer et al., 2014) to knock-in a fast-folding *Venus* reporter into the endogenous *etv2* locus and generated a stable *etv2^Gt(2A-Venus)ci48^* reporter line (further abbreviated as *etv2^Gt(2A-Venus)^*) (Figure S3A). *etv2^Gt(2A-Venus)+/-^* embryos displayed *Venus* expression in vascular endothelial progenitors and differentiated vascular endothelial cells, red blood cells and macrophages, similar to the previously established Gal4 knock-in line *etv2^Gt(2A-Gal4)^; UAS:GFP* (Chestnut and Sumanas, 2019) (Figure S3B-G). While heterozygous *etv2^Gt(2A-Venus)+/-^* embryos appeared normal, homozygous *etv2^Gt(2A-Venus)-/-^* embryos showed severe defects in vascular development, associated with the *etv2* loss-of-function phenotype. *etv2^Gt(2A-Venus)+/-^* embryos recapitulated endogenous *etv2* expression in SVF cells (Figure S3D inset), which has been challenging to observe in other previously established GFP reporter lines, likely due to the longer maturation time of GFP protein compared to *Venus*. FISH analysis confirmed co-localization of *etv2* mRNA and *Venus* fluorescence in the same SVF cells in *etv2^Gt(2A-Venus)+/-^* embryos (Figure S3H). We then performed time-lapse imaging from 24 to approximately 40 hpf using *etv2^Gt(2A-Venus)+/-^* crossed to *Tg(kdrl:mCherry)* reporter line which label vascular endothelial cells. During this early period, we observed multiple SVF cells which emerged in the mesenchyme adjacent to the PCV, and subsequently migrated and incorporated into the PCV (Figure 3A, white arrowheads, Videos S2 and S3). Some SVF cells proliferated during this process (Figure 3A, magenta and yellow arrowheads). New SVF progenitors continued to emerge during the imaging period.

**Figure 3.**
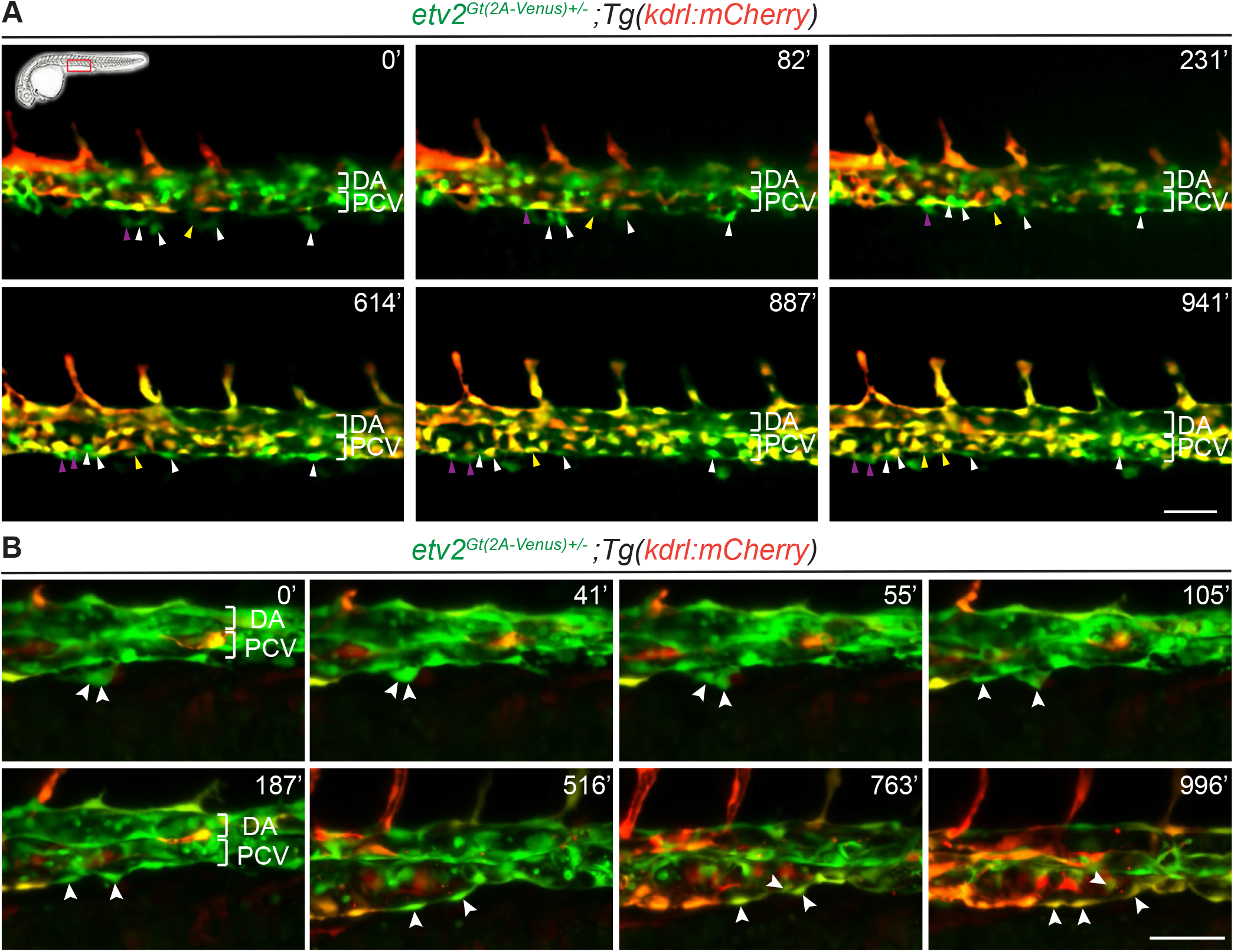
SVF cells contribute to axial vasculature and intercalate into the posterior cardinal vein: (A) Time-lapse images of *etv2^Gt(2A-Venus)+/-^* ; *Tg(kdrl:nls-mCherry)* embryo showing multiple SVF cells integrating into the PCV (white arrowheads) between 24 and 40 hpf. Images also show proliferation of SVF cells after integrating into PCV (magenta and yellow arrowheads). (B) High magnification time-lapse images of *etv2^Gt(2A-Venus)+/-^* ; *Tg(kdrl:mCherry)* embryo showing the sequential steps of SVF cell integration into the PCV (white arrowheads) between 24 and 40 hpf. An SVF cell contacts PCV, followed by cell rearrangement in the PCV, SVF cell proliferation and integration into the PCV. SVF cells initially express only *Venus* followed by *mCherry* expression upon integration and proliferation. Time is shown in minutes after 24 hpf. PCV, posterior cardinal vein; DA, dorsal aorta. Scale bars: A: 100 µm, B: 50 µm.

Higher magnification imaging showed that SVF cells made contact with VECs in the PCV prior to their integration. Subsequently, adjacent VECs in the PCV rearranged and appeared to make contact with SVF cells. Some SVF cells proliferated at this time. In the final stage, the SVF cells integrated into the PCV (Figure 3B, Video S4). While SVF cells initially had no or very little *kdrl:mCherry* expression, they upregulated *kdrl:mCherry* after integrating into the PCV which confirms their differentiation into vascular endothelial cells.

To follow SVF cell contribution during a longer time interval, we performed time-lapse imaging from 24 to approximately 70 hpf using the *etv2^Gt(2A-Venus)+/-^* crossed to *Tg(kdrl:nls-mCherry)* reporter line which expresses mCherry in the nuclei of vascular endothelial cells. During this period, we observed three different cell behaviors of the SVF cells. As described above, a subset of the SVF cells migrated and incorporated into the PCV (24%) (Figure S4A, white arrowheads, Videos S3, S5) by 48 hpf. However, many SVF cells contributed to the subintestinal vasculature, including the SIA (28%) (Figure S4A, yellow arrowheads) and SIV (48%) (Figure S4A,B, blue arrowheads, Video S5). While the SIA and SIV are known to be derived from the PCV (Hen et al., 2015), SVF cells appeared to incorporate directly into the SIA and SIV without first migrating to the PCV, and subsequently proliferated (Video S5, yellow and blue arrowheads). The progeny of each SVF cell contributed only to a single type of vessel (PCV, SIA or SIV, Table S1).

### SVF cells may form new VECs and contribute to vascular recovery

To examine the functional potential of SVF cells, we tested if SVF cells could form new VECs in the absence of axial vessels and contribute to vascular recovery. We utilized a nitroreductase (NTR) – metronidazole (MTZ) approach (Curado et al., 2008; Chlebowski et al., 2017) to ablate all early-forming endothelial cells except for SVF cells. In the presence of NTR, MTZ is metabolized to a toxic compound which induces apoptosis in the cells expressing NTR without affecting neighboring cells. We had previously generated an *etv2^Gt(2A-Gal4)ci32^* gene trap line (further abbreviated as *etv2^Gt(2A-Gal4)^*) which has an insertion of the Gal4 gene/transcriptional activator within the *etv2* coding sequence. As described previously, heterozygous *etv2^Gt(2A-Gal4)^* embryos recapitulate the endogenous expression pattern of *etv2* and do not show any significant defects in vascular development (Chestnut and Sumanas, 2019). The *etv2^Gt(2A-Gal4)^* line was crossed into an *UAS:GFP;UAS:mCherry-NTR* background which resulted in the expression of GFP, mCherry and NTR under the control of the *etv2* promoter (Figure 4A, E, I).

**Figure 4.**
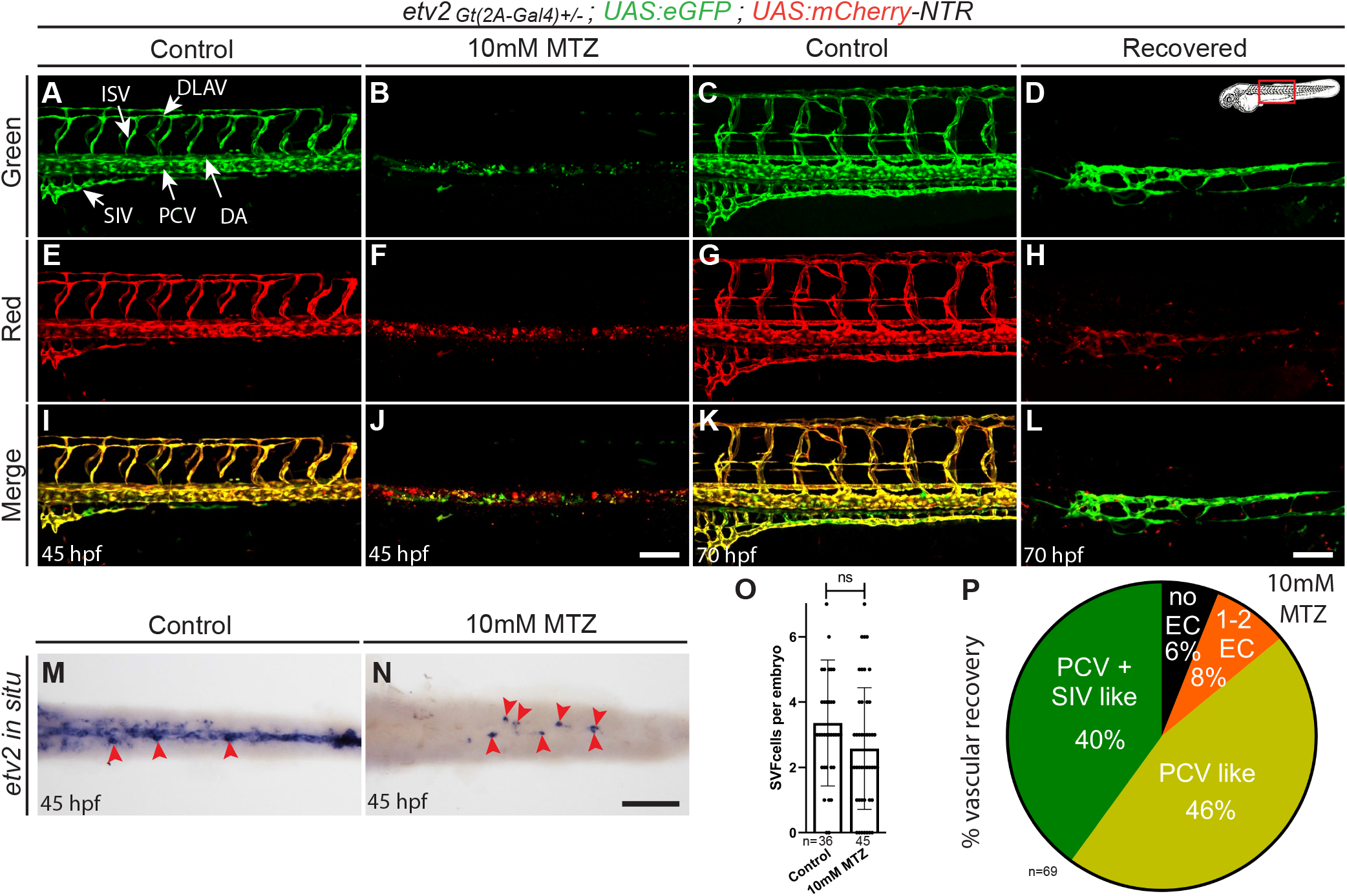
SVF cells contribute to vascular recovery after endothelial cell ablation: (A-L) *etv2^Gt(2A-Gal4)+/-^;UAS:GFP;UAS:mCherry-NTR* embryos were treated with MTZ from 50% epiboly to 45 hpf. Subsequently, MTZ was washed out and embryos were allowed to recover until 70 hpf. Note the complete ablation of *etv2* expressing endothelial cells in MTZ treated embryos at 45 hpf compared to 0.1% DMSO treated control embryos (A,B,E,F,I,J). Formation of new vascular cords at the position of the PCV and SIV was observed in the recovered embryos at 70 hpf (D,H,L). (M,N) Whole mount *in situ* hybridization (WISH) of embryos at 45 hpf showing expression of *etv2* only in SVF cells in MTZ treated embryos (red arrowheads). (N) quantification of SVF cells in MTZ treated (n=45) and control DMSO treated embryos (n=36). Mean ± s.d, ns: not significant (two-tailed, unpaired *t*-test). Each data point represents one embryo. (O) Percentage of embryos at 70 hpf showing recovered individual endothelial cells (EC) or partial blood vessels observed at the position of PCV and / or SIV (n=69). Data from 3 independent experiments. PCV, posterior cardinal vein; DA, dorsal aorta; ISV, inter-segmental vessels; DLAV, dorsal longitudinal anastomosing vessel; SIV, subintestinal vein; SIA, supraintestinal artery; EC, endothelial cell. Scale bars: 100 µm.

Because SVF cells emerge later than other VECs, we reasoned that early treatment with MTZ would ablate all VECs while SVF cells would be left intact. We treated embryos with 10 mM MTZ from 50% epiboly (∼5 hpf) to 45 hpf stages. At 45 hpf, embryos showed complete ablation of VECs expressing the *etv2* reporter (Figure 4A, B, E, F, I, J). This was confirmed by WISH analysis of embryos against *etv2*. Untreated embryos showed expression of *etv2* in the VECs as well as SVF cells. MTZ-treated embryos showed complete ablation of VECs, while SVF cells were still present bilaterally along the yolk extension (Figure 4M, N). Quantification of SVF cells revealed no significant difference between treated and untreated embryos (Figure 4O). MTZ was washed out at 45 hpf and embryos were analyzed for vascular recovery at 70 hpf. At this stage, we observed the emergence of VECs and formation of vascular tubes in the MTZ-treated embryos (Figure 4A, C, D, G, H, K, L and Video S6). 46% of embryos showed partially formed blood vessels at the anatomical position of the PCV, while 40% of embryos had partial vasculature at the positions of PCV and SIV. 8% of embryos showed GFP expression in only 1-2 individual VECs, while 6% embryos did not form any VECs (Figure 4P). Because only SVF cells showed *etv2* expression at 45 hpf after VEC ablation, these data suggest that recovered vessels are derived from SVF cells.

### *etv2* and *tal1* function is necessary for the recovery of vasculature

Both *etv2* and *tal1* are known critical regulators of vascular and hematopoietic development(Patterson et al., 2005; Sumanas and Lin, 2005). We therefore investigated their function in the recovery of vasculature in MTZ-treated embryos. To investigate the effect of the loss of *etv2* function, we used *etv2^Gt(2A-Gal4)-/-^; UAS:GFP;UAS:mCherry-NTR* embryos, which are deficient in *etv2* function and have severe vascular defects consistent with previous studies (Chestnut and Sumanas, 2019; Sumanas and Lin, 2005) (Figure 5A-D). We treated *etv2* mutant and heterozygous embryos with MTZ as described in the previous section. By 45 hpf, MTZ treatment resulted in complete ablation of VECs. After washing out MTZ and examining embryos at 70 hpf for VEC recovery (Figure 5E-H), we observed partially recovered vessels at the position of either PCV or SIV in only 26% of *etv2* homozygous embryos compared to 65% of *etv2* heterozygous embryos. Furthermore, 65% of *etv2^Gt(2A-Gal4)-/-^* embryos showed no recovery compared to 13% of *etv2^Gt(2A-Gal4)+/-^*embryos (Figure 5Q). These observations suggest that *etv2* function is essential for the differentiation of SVF cells and the recovery of vasculature.

**Figure 5.**
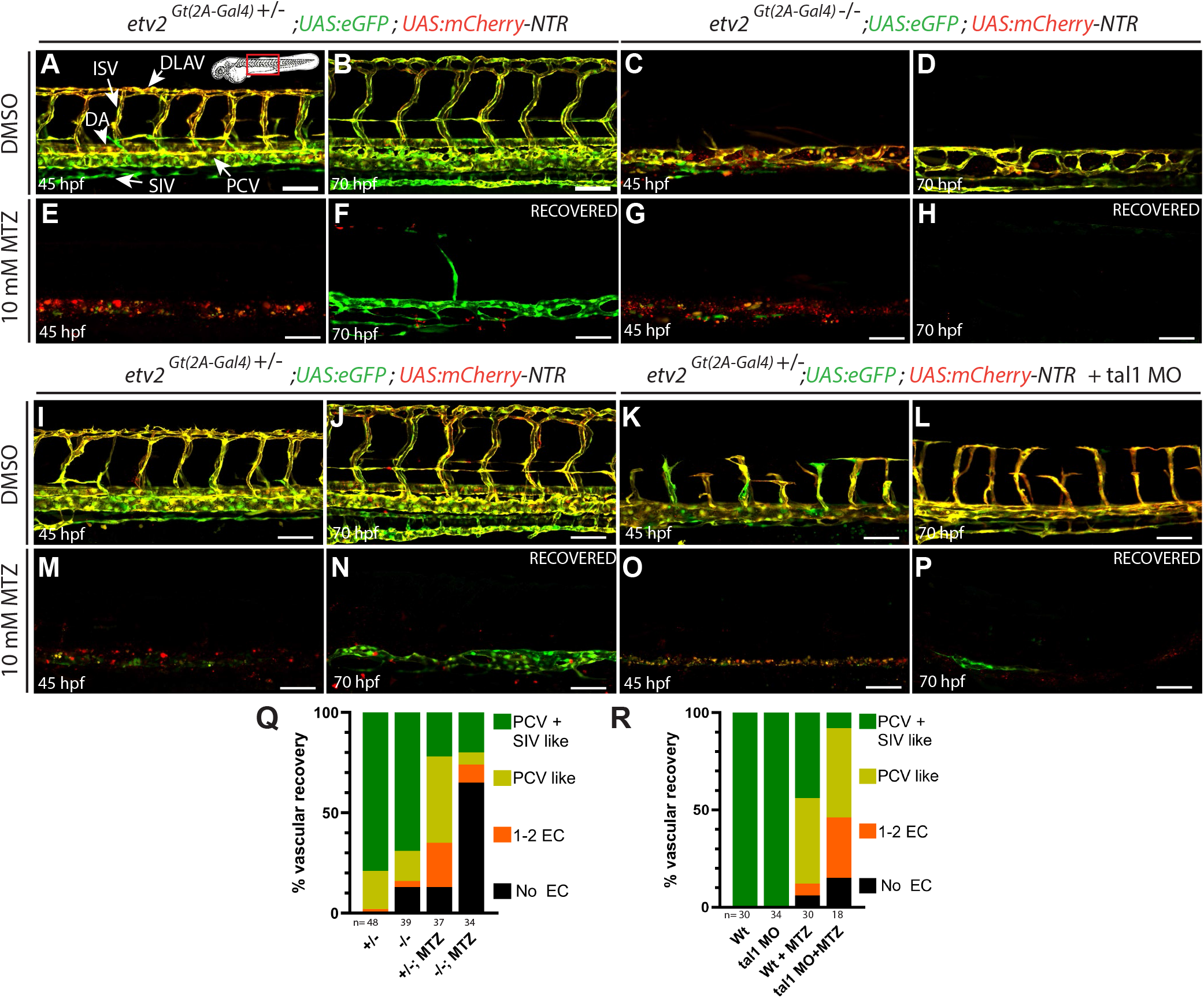
*etv2* and *tal1* function is necessary for recovery of vasculature after endothelial cell ablation: (A-H) Confocal images of *etv2^Gt(2A-Gal4)^;UAS:GFP;UAS:mCherry-NTR* homozygous mutant embryos showing severe vascular defects compared to heterozygous embryos at 45 and 70 hpf (A-D). *etv2^Gt(2A-Gal4)^;UAS:GFP;UAS:mCherry-NTR* embryos were treated with MTZ from 50% epiboly to 45 hpf to ablate *etv2* expressing endothelial cells. MTZ was washed out at 45 hpf and embryos were allowed to recover (E-H). *etv2* homozygous mutants show no or greatly reduced vascular recovery at 70 hpf compared to heterozygous sibling embryos. (Q) A percentage of embryos showing no recovery (no EC), only 1-2 ECs, or partially formed blood vessels at the anatomical position of PCV (PCV-like) or at the positions of both PCV and SIV (SIV-like). (I-P) Confocal images of *etv2^Gt(2A-Gal4)+/-^; UAS:GFP;UAS:mCherry-NTR* embryos injected with *tal1* MO showing severe vascular defects at 45 and 70 hpf (I-L). *etv2^Gt(2A-Gal4)+/-^ ;UAS:GFP;UAS:mCherry-NTR* embryos were injected with *tal1* MO at 1 cell stage and treated with MTZ from 50% epiboly to 45 hpf to ablate *etv2* expressing endothelial cells. MTZ was washed out at 45 hpf and embryos were allowed to recover. *tal1* morphants show significantly reduced vascular recovery (M-P). (R) A percentage of embryos showing no recovery (no EC), only 1-2 ECs, or partially formed blood vessels at the anatomical position of PCV (PCV-like) or at the positions of both PCV and SIV (SIV-like). Data from 2 independent experiments. PCV, posterior cardinal vein; SIV, subintestinal vein; EC, endothelial cell. Scale bars: 100 µm.

To evaluate the function of *tal1* in vascular recovery, we used a splice blocking *tal1* morpholino (MO) which has been previously validated (Dooley et al., 2005). *etv2^Gt(2A-Gal4)+/-^;UAS:GFP;UAS:mCherry-NTR* embryos were injected with *tal1* MO at the 1 cell stage and treated with MTZ from 50% epiboly to 45 hpf stages. As reported in previous studies (Dooley et al., 2005; Patterson et al., 2005), *tal1* MO-injected embryos (morphants) showed vascular defects (Figure 5I-P). Subsequently, MTZ was washed out at 45 hpf and embryos were allowed to recover till 70 hpf (Figure 5M-P). 54% of *tal1* morphants showed partially recovered vessels at the position of the PCV or SIV compared to 88% of control embryos (Figure 5R), suggesting that *tal1* function is necessary for the formation of SVF cells and recovery of vasculature.

### Transcriptome analysis of SVF cells reveals their unique transcriptional profile

To identify the transcriptional signature of SVF cells, we analyzed the transcriptome dataset previously obtained by single-cell RNA-seq (scRNAseq) approach of cells isolated from zebrafish embryo trunks dissected at 30 hpf (Metikala et al., 2021). In this study, over 20,000 cells were disaggregated from dissected zebrafish embryo trunks and subjected to RNA-seq resulting in the identification of transcriptional profiles for 27 different cell types. However, only a single population of vascular endothelial cells was identified following the initial cell clustering (Figure S5A), and no candidate SVF cell cluster was observed. We reasoned that SVF cells comprise only a small fraction of all trunk cells, and therefore they may not form a separate cluster using standard analysis approaches. Therefore, we filtered the normalized gene expression matrix for high expression of SVF marker genes *etv2, tal1,* and *lmo2* (log_2_>1). Based on our *in situ* hybridization results, SVF cells are the only cells in the trunk region that show significant co-expression of all three marker genes. We also excluded cells with significant expression of *fli1a,* which is expressed in differentiated VECs but has no or low expression in SVF cells (Figure S2). Such approach identified 23 candidate SVF cells (Figure S5). Differential expression analysis identified over 30 additional genes preferentially enriched in this candidate SVF cell population, which included known regulators of vascular development *sox7* (Hermkens et al., 2015)*, egfl7* (Parker et al., 2004)*, tmem88a* (Cannon et al., 2013) as well as many genes which have not been previously associated with vascular development (Table S2). To validate these results, we performed WISH on selected marker genes. Indeed, expression of *tmem88a, sox7* and *egfl7* was observed in SVF cells in addition to their previously described expression in blood vessels (Cannon et al., 2013; Hermkens et al., 2015; Parker et al., 2004) (Figure S6). We have recently described a transcriptional profile of vascular endothelial progenitor cells (EPCs) using scRNA-seq performed at an earlier stage of 20-somites in zebrafish embryos (Chestnut et al., 2020). The transcriptional profile of SVF cells partially overlapped with the transcriptional profile of EPCs, and *etv2, tal1* and *lmo2* were among the top marker genes in both cell populations (Table S3). However, some marker genes including *jam2b* and *nanos1* were unique to SVF cells and were absent from or had a very low expression in EPCs, while others including *robo3* and *dusp5* had high expression in EPCs and low expression in SVF cells. These data suggest that SVF cells have a unique transcriptional profile which is distinct from endothelial progenitor cells that are present during early vasculogenesis.

### *etv2* and *tal1* function is required for differentiation of SVF cells

We hypothesized that *etv2* and *tal1* may function upstream of other factors during the differentiation of SVF cells. To study the role of *etv2*, we performed WISH analysis for *tal1*, *tmem88a, sox7* and *egfl7* expression in *etv2^y11^* mutants (Pham et al., 2007). The number of SVF cells labeled by these markers was greatly reduced in *etv2* mutants, and the expression of these genes in the remaining SVF cells also appeared to be decreased (Figure 6A,C). In addition, the expression of *sox7* and *egfl7* was also reduced in endothelial cells. To examine if only the expression of these markers was reduced in SVF cells or if there was a general loss of SVF cells, we analyzed *etv2* mRNA expression in embryos injected with a combination of two previously validated *etv2* translation-blocking morpholinos (Sumanas and Lin, 2005). As previously reported, the *etv2* MO injection recapitulated severe vascular defects observed in *etv2* mutant embryos (Sumanas and Lin, 2005) (Figure S7A, C). A slight reduction in SVF cell number was observed in *etv2* morphants at 30 hpf (10.36 ± 3.28 SVF cells/embryo) compared to uninjected controls (12.32 ± 3.17 SVF cells/embryo) (Figure S7B,D,E). Because, as reported previously, *etv2* mRNA expression is strongly increased in *etv2* morphants, possibly due to a negative feedback loop (Sumanas and Lin, 2005; Sumanas et al., 2008), it is more challenging to identify SVF cells in morphants due to strong *etv2* expression in the adjacent vasculature. Therefore, it is possible that their numbers were slightly undercounted in *etv2* morphants. Regardless, it was evident that there were more *etv2*-expressing SVF cells than the cells that express other SVF markers in *etv2* inhibited embryos. This argues that *etv2* functions upstream of *tal1, tmem88a, sox7 and egfl7* expression and SVF cells fail to differentiate in *etv2-*inhibited embryos.

**Figure 6.**
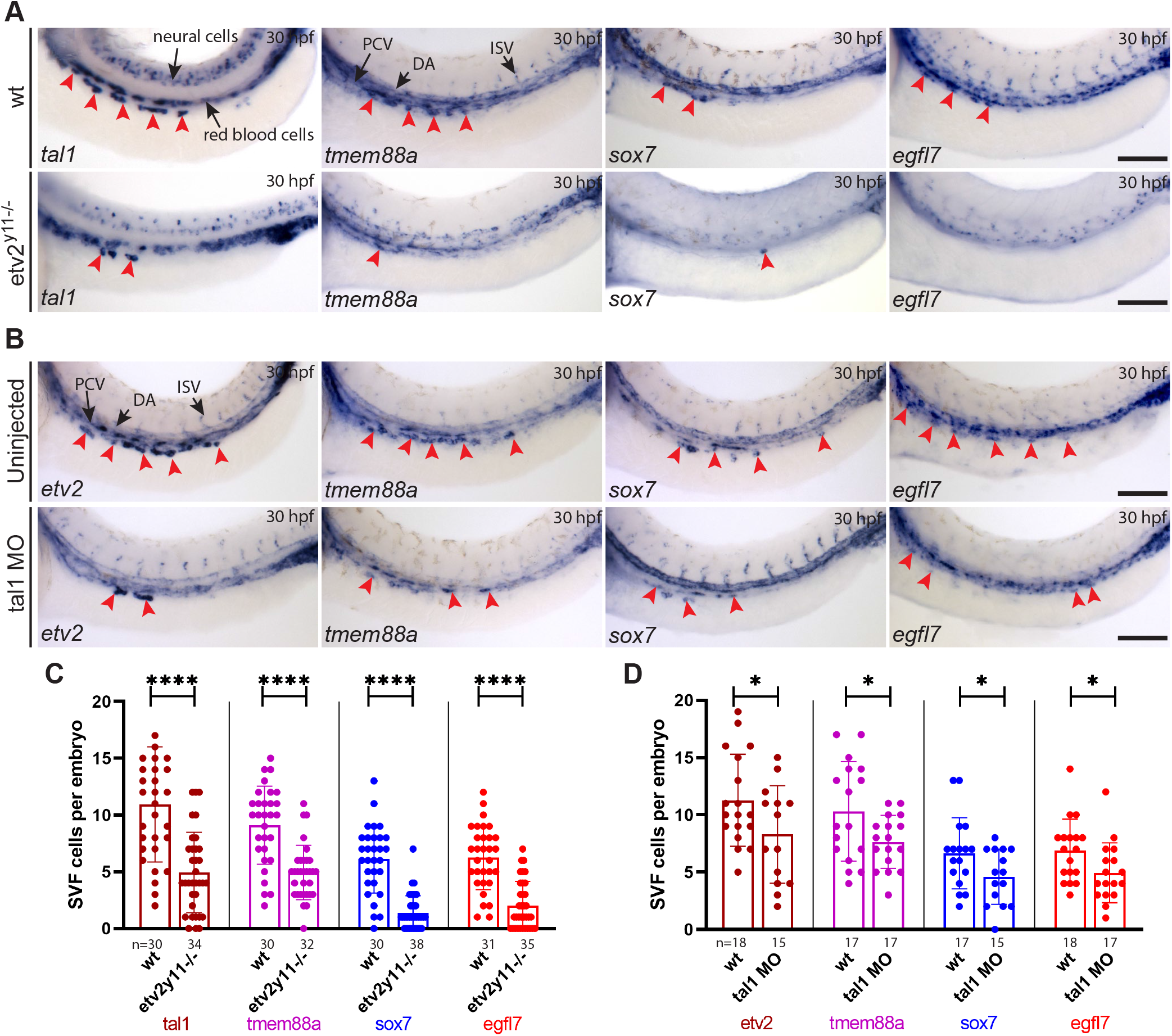
*etv2* and *tal1* function is necessary for *tmem88a*, *sox7* and *egfl7* expression in SVF cells: (A) WISH images of *etv2^y11-/-^* mutants and wt siblings at 30 hpf showing SVF cells (red arrowheads). WISH was performed using *tal1*, *tmem88a*, *sox7* and *egfl7* probes. (B) WISH images of *tal1* morphants and wt siblings at 30 hpf showing SVF cells (red arrowheads). WISH was performed using *etv2*, *tmem88a*, *sox7* and *egfl7* probes. (C) Quantification of SVF cells in *etv2 ^y11-/-^* mutants and wt siblings. Only wild-type (+/+) and null mutants (**-**/**-**) were used for analysis. (D) Quantification of SVF cells in *tal1* MO-injected embryos and uninjected (UI) controls (*etv2*: UI, n=18, *tal1* MO n=15; *tmem88a*: UI, n=17, *tal1* MO n=17; *sox7*: UI, n=17, *tal1* MO n=15; *egfl7*: UI, n=18, *tal1* MO n=17). mean ± s.d, \**P*<0.05, \*\*\*\**P*<0.0001 (two-tailed unpaired t-test). Each data point represents one embryo. Data from 2 independent experiments. UI, un-injected; PCV, posterior cardinal vein; DA, dorsal aorta; ISV, intersegmental vessels. Scale bars: 100 µm.

To study the role of *tal1* in SVF cell formation, we injected embryos with a previously validated splice-blocking *tal1* morpholino (Dooley et al., 2005). The number of SVF cells that express *etv2*, *tmem88a, sox7* and *egfl7* was reduced significantly in *tal1*-deficient embryos (Figure 6B,D). However, expression of these markers was not significantly reduced in endothelial cells. Collectively, these results show that *etv2* and *tal1* function is necessary for the expression of *tmem88a, sox7* and *egfl7* in SVF cells.

### Loss of SVF cells results in defective subintestinal vasculature

We then aimed to interrogate the functional role of SVF cells during normal development. Although currently available tools do not allow specific ablation of the SVF cells, we reasoned that inducible *etv2* inhibition after initial vasculogenesis is complete would inhibit the differentiation of SVF cells. We have previously demonstrated that a photoactivatable *etv2* MO can be used for specific inhibition of *etv2* function at different time points (Kohli et al., 2013). Photoactivation of this *etv2* MO at early stages (6 hpf) phenocopied the *etv2* mutant phenotype, while embryos injected with the photoactivatable *etv2* MO did not show significant vascular defects at 22 hpf in the absence of photoactivation (Figure S8A-C). To determine if *etv2* inhibition affected the differentiation of SVF-derived endothelial cells, we injected this photoactivatable *etv2* MO into *Tg(fli1a:kaede)* embryos followed by photoactivation at 22 hpf and Kaede photoconversion at 24 hpf. We then analyzed the formation of new VECs and defects in vascular patterning. *etv2* morphants showed defects in the formation of the SIA (Figure 7A,B, white arrowheads). 18% of analyzed embryos exhibited an absence of the SIA, while 61% of embryos showed only partial SIA formation. In addition, a subset of embryos showed minor defects in the formation of other blood vessels in the trunk. *Etv2* morphants also showed significantly reduced expression of *tal1* in SVF cells at 30 hpf compared to uninjected embryos (Figure 7C), arguing that inducible inhibition of *etv2* function effectively inhibited differentiation of SVF cells. Moreover, quantification of new VECs showed that there were fewer green-only new VECs in the SIA of *etv2* morphants (Figure 7A,D, magenta arrowheads). Embryos injected with the morpholino and not subjected to photoactivation did not show any significant vascular defects (Figure S8D, E, G). To confirm these results, we analyzed genetic *etv2^y11^* mutant embryos (Pham et al., 2007). As reported previously, *etv2* mutant embryos showed strong defects in vasculogenesis and angiogenesis, which reflects a known *etv2* requirement for early vascular development. Intriguingly, PCV and SIV were present although they were dysmorphic and did not lumenize (Figure S8F). In contrast, SIA could not be located in *etv2^y11^* mutant embryos, which confirms the phenotype observed in embryos injected with photoactivatable etv2 MO. Altogether, these observations argue that *etv2* inhibition at 22 hpf results in defective SIA development and causes reduced numbers of new VECs within the subintestinal vasculature. Because our data show that the SVF is the source of many (if not all) new VECs in the trunk region after 24 hpf, these results suggest that *etv2* function in SVF cells is required for SIA formation.

**Figure 7.**
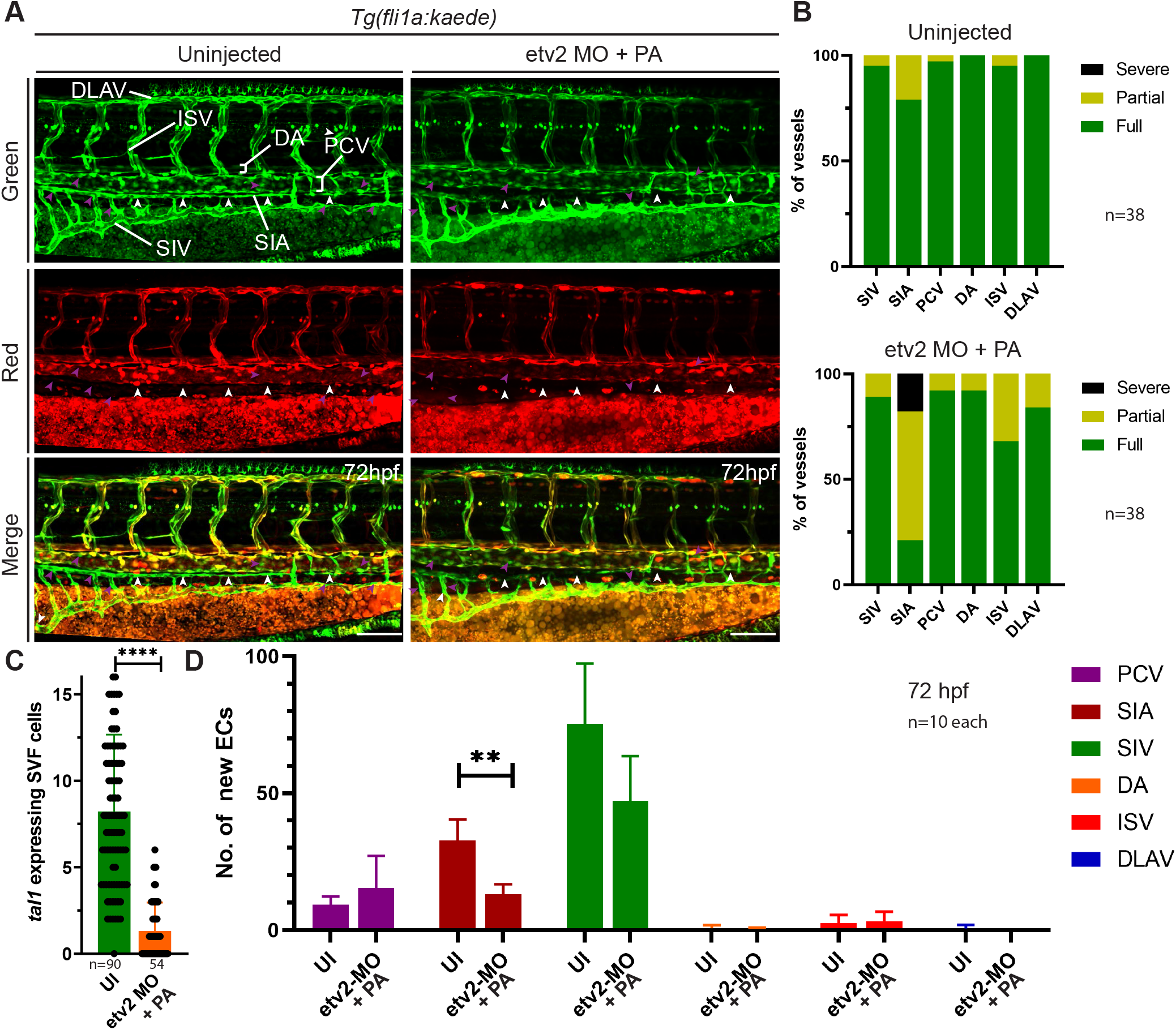
Loss of SVF cells results in defective subintestinal vasculature: (A) Photoactivatable *etv2* MO was injected in *Tg(fli1a:Kaede)* embryos followed by *etv2* MO photoactivation at 22 hpf and *Kaede* photoconversion at 24 hpf. Embryos at 72 hpf show defective SIA formation (white arrowheads) and the presence of new VECs (magenta arrowheads). (B) Quantification of fully formed, partially formed or absent vessels in the trunk in control uninjected embryos (n=38) and *etv2* morphants (n=38). (C) Quantification of *tal1* expressing SVF cells at 30 hpf in *etv2* morphants (n=54) and wt siblings (n=90). Each data point represents one embryo. (D) Quantification of new VECs in *etv2* morphants (n=10) and wt siblings (n=10) at 72 hpf. mean ± s.d, \*\**P*=0.001, \*\*\*\**P*<0.0001 (two-tailed unpaired t-test). Data from 4 independent experiments. PCV, posterior cardinal vein; DA, dorsal aorta; ISV, intersegmental vessels; DLAV, dorsal longitudinal anastomosing vessel; SIV, subintestinal vein; SIA, supraintestinal artery. Scale bars: 100 µm.

### *VEGF* signaling and *npas4l* function is necessary for the formation of SVF cells

Several signaling pathways such as *wnt* (Gore et al., 2011; Hübner et al., 2017), *bmp* (Lee et al., 2008; Walmsley, 2002) and *vegf* (Carmeliet et al., 1996; Ferrara et al., 1996) have been previously implicated in blood vessel formation. To investigate the role of *vegf* signaling in the formation of SVF cells, we first analyzed *etv2* expression in SVF cells in *vegfaa* and *kdrl* mutants, which are considered zebrafish homologs of mammalian *Vegfa* and its receptor *Vegfr2 / Flk1,* respectively. We observed a significant reduction in the number of SVF cells in both *vegfaa^-/-^* and *kdrl^-/-^* mutants at 30 hpf compared to their respective wild-type siblings, (Figure 8A, B, D, E) suggesting that *vegf-kdrl* pathway is required for the formation of SVF cells. To confirm the requirement of *vegf* signaling for SVF cell differentiation, we treated embryos using pan-*vegfr* inhibitor SU5416 (Serbedzija et al., 1999) and injected a previously validated *vegfaa* morpholino (Nasevicius et al., 2000). Both resulted in a significant reduction in the number of SVF cells (Figure S9 A, B, D). These results suggest that *vegf-kdrl* pathway is required for the formation of SVF cells.

**Figure 8.**
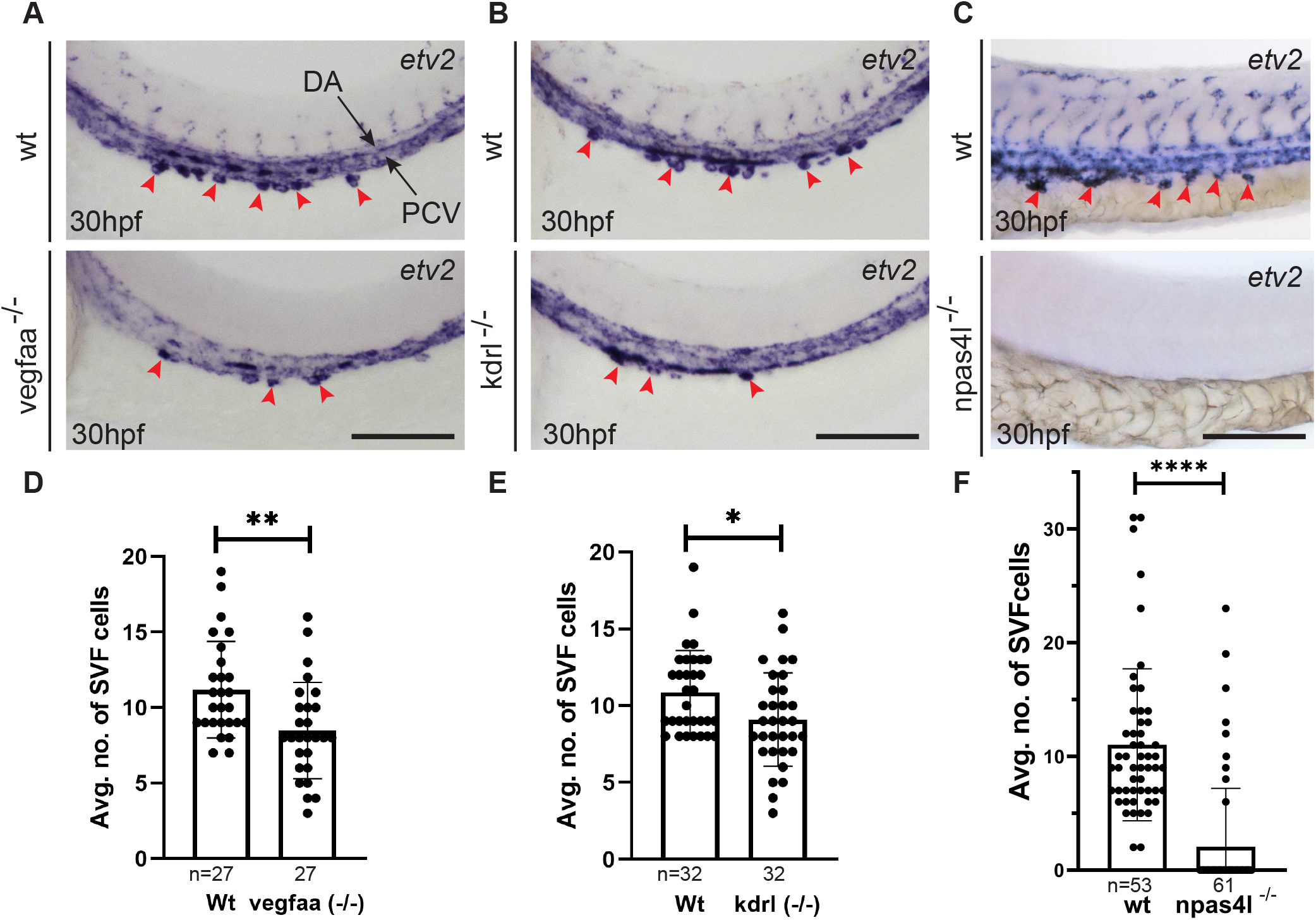
*Vegf* signaling pathway and *npas4l* function is necessary for the formation of SVF cells: (A) Whole mount *in situ* hybridization (WISH) analysis for *etv2* expression at 30 hpf in *vegfaa^-/-^* mutants and their wild-type siblings. (B) WISH analysis for *etv2* expression at 30 hpf in *kdrl^-/-^* mutants and their wild-type siblings. (C) WISH analysis for *etv2* expression at 30 hpf in *npas4l^-/-^* mutants and their wild-type siblings. SVF cells are shown using red arrowheads. (D) Quantification of SVF cells in wild-type (n=27) and *vegfaa^-/-^* mutants (n=27) embryos. (E) Quantification of SVF cells in wild-type (n=32) and *kdrl^-/-^* mutants (n=32) embryos. (F) Quantification of SVF cells in wild-type siblings (n=53) and *npas4l^-/-^* embryos (n=61). *vegfaa^-/-^* and *kdrl^-/-^* mutants were analyzed in 2 independent experiments and *npas4l^-/-^* in 3 independent experiments. Note that wild-type siblings include a mixture of wild-type and heterozygous embryos in all experiments. mean ± s.d, \**P*<0.05, \*\**P*<0.005 and \*\*\*\**P*<0.0001 (two-tailed unpaired t-test). Each data point represents one embryo. PCV, posterior cardinal vein; DA, dorsal aorta. Scale bars: 100 µm.

Subsequently, we treated zebrafish embryos starting at 50% epiboly with chemical inhibitors for *wnt* and *bmp* pathways and analyzed *etv2* expression in SVF cells at 28 hpf. Inhibition of *wnt* and *bmp* signaling did not significantly affect the number of SVF cells (Figure S9C, E) suggesting these pathways are not necessary for SVF cell formation.

Furthermore, *cloche/npas4l* has been shown to play a crucial role in vasculogenesis. *npas4l^-/-^* mutants show severe defects in vascular network formation (Reischauer et al., 2016; Stainier et al., 1995). Analysis of *etv2* expression by WISH showed that SVF cells were completely absent or greatly reduced in *cloche* mutants (Figure 8C,F). These observations suggest that *npas4l* functions upstream of *etv2* during SVF cell formation.

### Identification of SVF-like cells in murine embryos

It is currently unclear if SVF-like cells are present in mammalian embryos. Intriguingly, Etv2-Venus positive cells that were negative for VE-cadherin have been previously reported outside of the established vasculature in the head and trunk region of murine embryos at E8.5 and E9.5 (Kobayashi et al., 2013). To examine these cells in greater detail, we analyzed Etv2-Venus reporter embryos at E10.5. Etv2-Venus positive cells negative for ERG and PECAM expression were observed adjacent to mesonephros outside of the cardinal vein (CV) (Figure 9A-D, yellow arrowheads). Some Etv2-Venus cells were located in the CV, and were positive for ERG and negative for PECAM, suggesting that they had recently incorporated into the CV (Figure 9A, E-G yellow arrowheads). To perform lineage tracing of late-forming Etv2+ cells, we mated *Etv2^T2A-iCreERT2/+^* knock-in and *ROSA^tdTomato/tdTomato^* reporter mice and injected tamoxifen at E9.5. *Etv2^T2A-CreERT2/+^; ROSA^tdTomato/+^* embryos were isolated at E10.5 and stained to detect ERG, a transcription factor expressed in vascular endothelial cells, and tdTomato to detect Etv2^+^ cells and their lineage. A subset of endothelial cells in the CV and intersegmental vessels expressed tdTomato (Figure 9H). This suggests that late forming Etv2+ progenitors contribute to the CV and ISVs in mouse embryos. Intriguingly, ERG^-^, tdTomato^+^ cells were observed adjacent to the CV in the vicinity of the mesonephros. These cells appeared more round compared with differentiated vascular endothelial cells and did not express ERG (Figure 9H-K). Together, these data show that Etv2^+^ progenitors negative for the expression of differentiated vascular markers are also present in the trunk region of murine embryos outside of the established vasculature, suggesting that SVF cells are evolutionarily conserved between zebrafish and mammalian embryos.

**Figure 9.**
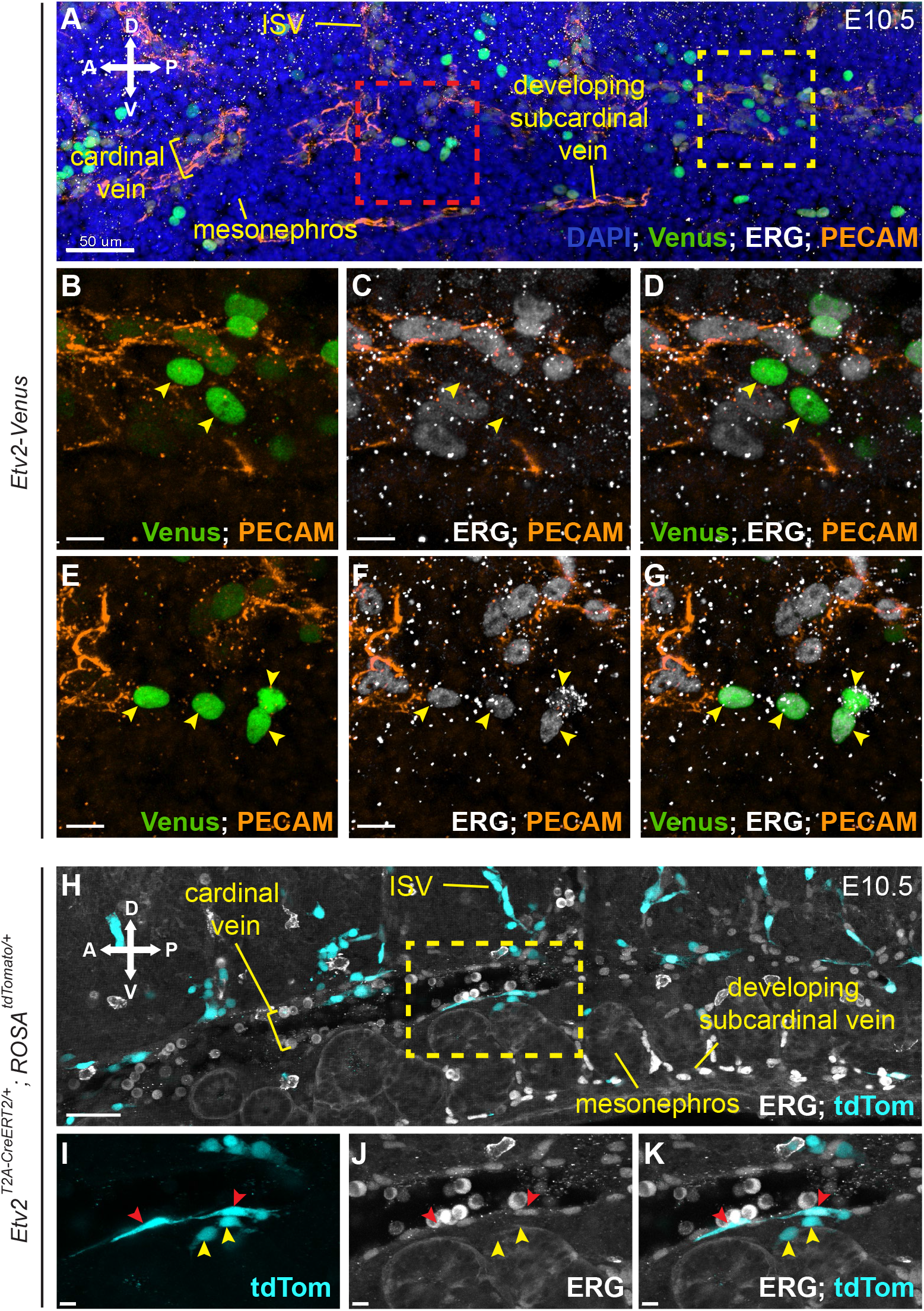
Undifferentiated Etv2+ progenitors are present outside of the existing vasculature in the trunk region of murine embryos: (A-G) Etv2-Venus reporter embryo at E10.5 stained to detect Venus (green), ERG (white) and PECAM (orange). (A) Sagittal view of 15 µm z-stack through the cardinal vein and developing subcardinal vein of a 28s mouse embryo showing Etv2-Venus positive cells (green) adjacent to mesonephros outside of the cardinal vein. (B-D) Corresponding expanded view (yellow box, A) of two Etv2-Venus positive cells that are negative for ERG and PECAM (yellow arrowheads, C). (E-G) Corresponding expanded view (red box, A) of four Etv2-Venus positive cells (green) that are positive for ERG (white) but negative for PECAM (orange) (yellow arrowheads, G). (H-K) Etv2T2A-CreERT2/+ males were mated with homozygous Ai9 reporter females and treated with tamoxifen at E9.5. Embryos were collected at E10.5 and stained to detect ERG (white) and tdTomato (turquoise). (H) Sagittal view of 15 µm z-stack through the cardinal vein and developing subcardinal vein of a 35s mouse embryo. (I-K) Corresponding expanded view (yellow box, H) of two tdTomato positive cells, (yellow arrowheads) directly ventral to the cardinal vein that are ERG negative. tdTomato positive cells which are incorporated into existing vasculature express ERG (red arrowheads, I-K).

## Discussion

It has been generally thought that vascular progenitors give rise to the formation of new blood vessels by the mechanism of vasculogenesis, while new vascular growth after the formation of the initial vasculature occurs largely by angiogenic sprouting. Although there have been previous reports of endothelial progenitor cells in adults that may contribute to the vascular growth or regeneration (Asahara et al., 1997; Shi, et al., 1998), this has been debated (Yoder, 2012). Our study demonstrates that new vascular progenitors emerge and contribute to functional embryonic vasculature after it has already initiated circulation. To our knowledge, this is one of the first demonstrations of the contribution of vascular progenitors to established vessels *in vivo*. A recent study has demonstrated that erythro-myeloid progenitors (EMP) can give rise to vascular endothelial cells and contribute to vascular growth in murine embryos (Plein et al., 2018). At this point, we do not have any evidence that SVF cells can contribute to erythroid or myeloid lineages, and their molecular signature matches the signature of vascular endothelial progenitors and not EMP cells. Therefore, incorporation of SVF cells into vasculature appears to be a distinct mechanism from the previously demonstrated vascular contribution of EMPs.

It is intriguing that SVF cells appear bilaterally along the region of the yolk sac extension. Some cells were positioned laterally adjacent to the pronephros, while others were observed ventrally and medially. It is likely that all SVF cells may originate laterally to the pronephros, and subsequently migrate ventrally and medially before integrating into the PCV.

Single-cell RNA-seq analysis has enabled the identification of the transcriptional signature of SVF cells. Intriguingly, it is quite similar to the transcriptional profile of vascular endothelial progenitors, which we have recently identified at an earlier stage of 20-somite stage embryos (Chestnut et al., 2020). Both cell types share many of the key markers including *etv2, tal1, lmo2, tmem88a, sox7* and *egfl7*. Yet despite these similarities, SVF transcriptional profile is distinct from early EPCs and shows expression of some unique markers including *jam2b* and *nanos1*. This argues that SVF cells are late-forming vascular progenitors which are distinct from early EPCs. Although some SVF markers (*etv2*, *tal1* and *lmo2*) are also known to label hematopoietic progenitors (Robb et al., 1995; Thompson et al., 1998; Wareing et al., 2012), we have not observed any contribution of SVF cells to blood lineages. In addition, no blood cells that are green-only were observed by *fli1:Kaede* or *etv2:Kaede* photoconversion at 24 hpf, further arguing that SVF cells do not give rise to blood cells.

Our results show three major SVF cell behaviors and fate choice patterns: 1) integration into the PCV, 2) direct contribution to the SIV, 3) direct contribution to the SIA (Figure S10). The PCV is known to give rise to several different subtypes of endothelial cells including lymphatic vasculature and subintestinal vessels (Hen et al., 2015; Koenig et al., 2016; Nicenboim et al., 2015). Intriguingly, many SVF cells appear to contribute directly to subintestinal vasculature without incorporating into the PCV first. This suggests that subintestinal vasculature is derived from two different sources of progenitors, some of which are located in the PCV, while others are SVF progenitors that directly contribute to the subintestinal vasculature. Furthermore, both SVF and non-SVF cells contribute to subintestinal vasculature, suggesting that these cells can contribute to different endothelial subtypes and are not predetermined. One caveat is that *Kaede* photoconversion and time-lapse imaging was performed starting at 24 hpf, while some SVF cells may be present at even earlier stages. It is also currently unclear if SVF cells can contribute to lymphatic progenitors, which are known to be derived from the PCV.

*Etv2* has been previously demonstrated to be the major regulator of vascular endothelial differentiation and functions upstream of *tal1* and most other vascular markers (Sumanas and Lin, 2005). Thus, it is not surprising that expression of *tal1, tmem88a* and other SVF markers is absent or reduced in *etv2* mutant embryos. Intriguingly, based on *etv2* expression analysis in *etv2* morphants, SVF cells still form in the same region but fail to differentiate. Inhibition of *etv2* function starting at 24 hpf using photoactivatable MO results in major defects in SIA formation, which is consistent with the major contribution of SVF cells to the SIA. Although SIV formation was not strongly affected, there are several factors that may compensate for the absence of *etv2* function. ETS transcription factors show a high degree of redundancy, and we and others have previously demonstrated partial redundancy between *etv2, fli1b* and *ets1* during embryonic vasculogenesis and angiogenesis (Chetty and Sumanas, 2020; Craig and Sumanas, 2016; Pham et al., 2007). Inhibition of *etv2* function may inhibit SVF cell differentiation only partially, and cells may compensate through increased proliferation, resulting in only partial defects in subintestinal vasculature formation.

In this study we aimed to interrogate the functional role of SVF cells. However, in the absence of a marker that is specific to SVF cells, it is not possible to specifically ablate this cell population. Therefore, we utilized an NTR-MTZ approach to ablate all other endothelial cells that form earlier while SVF cells were left intact. >90% of embryos showed partial recovery of trunk vasculature after MTZ was washed out. Because *etv2* expression was only observed in SVF cells after MTZ treatment, we argue that the recovered vasculature originated from SVF cells. Intriguingly, the recovered vessels were positioned at the anatomical locations of the PCV and SIV, which is similar to the observed contribution of SVF cells to the vasculature. Vascular recovery was dependent on *etv2* and *tal1* function, which further supports the role of these genes in SVF cell differentiation.

Molecular pathways that regulate the emergence of SVF cells are currently unclear. Our study implicates *vegf* signaling in this process. Our previous work has demonstrated that *vegf* signaling is required to upregulate *etv2* expression although it is dispensable for its initiation (Casie Chetty et al., 2017). We demonstrate in this study that zebrafish *vegfaa* mutant and morpholino inhibited embryos show reduced number of *etv2-*positive SVF cells. Morpholino-injected embryos showed a stronger reduction in SVF cell number compared with *vegfaa* mutant embryos which could be explained by the genetic compensation apparent in *vegfaa* mutants. *vegfaa* mutant embryos are know to upregulate expression of *vegfab,* a related zebrafish homolog of mammalian *Vegfa* (El-Brolosy et al., 2019). Intriguingly, *vegfaa* is expressed in the endoderm close to the area where subintestinal vessels form during 28-48 hpf stages (Koenig et al., 2016). Although SVF cells do not have strong expression of *Vegf* receptors, interrogation of their transcriptional profile shows small yet detectable expression of *kdrl* (although no *kdr* or *flt4* expression). Indeed, *kdrl* mutants show significant reduction in SVF cell number. This suggests that SVF cells could directly respond to *vegf* signaling which then would upregulate *etv2* and *kdrl* expression as a part of a positive feedback loop. *Vegfaa-kdrl* pathway may regulate specification, differentiation and / or proliferation of SVF cells which remains to be determined. It is likely that additional yet to be identified signaling pathways are involved in the specification of SVF cells in this region.

Perivascular *etv2-*expressing cells have been previously reported in the head and trunk regions of mouse embryos at E8.5 and E9.5 (Kobayashi et al., 2013), suggesting that SVF-equivalent cells may be also present in mammalian embryos. Indeed, our results demonstrate that Etv2+ and ERG-negative cells are present outside of the CV in murine embryos at E9.5-10.5. They are located in the trunk region adjacent to mesonephros and appear remarkably similar to SVF cells in zebrafish embryos. Therefore, it is likely that late-forming Etv2+ cells also contribute to the mammalian vasculature. This is suggested by tdTomato-labeled cells within the murine CV and intersegmental vessels. While it is likely that many labeled cells are derived from the newly formed Etv2+ progenitors, at present we cannot exclude the possibility that some of the labeling may be due to residual Etv2-CreERT2 expression within the CV. Nevertheless, these results strongly suggest that SVF cells and the mechanism of their incorporation into established vasculature are evolutionarily conserved.

In summary, our study has identified a population of late forming vascular progenitor cells which contribute to blood vessels by a novel mechanism of integrating into established vasculature. These findings will be important in understanding mechanisms of vascular development and may promote development of novel therapies aimed at vascular repair and regeneration.

## Acknowledgements

We thank Andrew Koenig for the initial analysis of SVF cells, Matthew Kofron and Evan Meyer for their technical assistance during image acquisition and analysis (Confocal Imaging Core – CCHMC), Andrew Potter and Steve Potter for single cell dissociation protocol and assistance and Shawn Smith and Kelly Rangel (Gene expression core – CCHMC) for assistance in processing samples for scRNAseq, Jenna Oberstaller and Swamy R. Adapa for scRNAseq analysis advice and Joshua Waxman for kind donation of pCS2+_Kaede plasmid. This research was supported by grants from American Heart Association post-doctoral fellowship to Sanjeeva Metikala (19POST34400016) and National Institutes of Health (R21AI128445) and CCHMC RIP awards to Saulius Sumanas.

## Author contributions

S.M. performed most experiments in the study, generated *Tg(fli1a:Kaede)* line, analyzed data and wrote the manuscript, B.C. generated *etv2^Gt(2A-Venus)^* line, S.C.C. analyzed 30 hpf trunk scRNAseq data and edited the manuscript, E.P and O.N contributed to NTR-MTZ experiments. M.W. performed *Etv2* expression and lineage analysis of mouse embryos, S.A. supervised mouse experiments. S.S. conceived and supervised the project and edited the manuscript.

## STAR Methods

### Zebrafish lines

Zebrafish lines used in the study are as follows: wild-type AB, *TgBAC(etv2:kaede)^ci6^* (Kohli et al., 2013), *etv2^Gt(2A-Venus)ci48^* (further abbreviated as *etv2^Gt(2A-Venus)^*), *etv2^Gt(2A-Gal4)ci32^;UAS:GFP;UAS:mCherry-NTR*(Chestnut and Sumanas, 2019) (further abbreviated as *etv2^Gt(2A-Gal4)^;UAS:GFP;UAS:mCherry-NTR*), *Tg(kdrl:nls-mCherry)^is4^* (Wang et al., 2010), *Tg(kdrl:mCherry)^ci5^* (Proulx et al., 2010), *etv2^y11^* (Pham et al., 2007), *Tg(fli1a:GFP)^y1^* (Lawson and Weinstein, 2002a), *Tg(enpep:GFP)* (Seiler and Pack, 2011), *vegfaa^+/-^ ; Tg(flk1:mCherry)* (Rossi et al., 2016) and *kdrl^+/-^ ; Tg(fli1a:GFP)* (Covassin et al., 2006). To generate *Tg(fli1a:kaede)^ci49^*, Kaede ORF was amplified from pCS2+_Kaede plasmid (kindly donated by J. Waxman) with oligos containing attb sequences (Kaede_attb_F5’-GGGGACAAGTTTGTACAAAAAAGCAGGCTTCACCatggtgagtctgattaaaccag, Kaede_attb_R5’–GGGGACCACTTTGTACAAGAAAGCTGGGTTttacttgacgttgtccggca, Q5 DNA polymerase (NEB, M0491S), Tm=72°C, amplicon length = 681 bp) and cloned into pDONR 221 using Gateway™ BP Clonase™ II Enzyme mix (Invitrogen, Cat. No: 11789020). Kaede ORF, fli1EP and polyA sequences from pDONR221_Kaede, p5Efli1ep (Villefranc et al., 2007) and p3E_polyA (Villefranc et al., 2007) respectively were cloned into pDEST-Tol2PA2 vector using LR Clonase™ II Plus enzyme (Invitrogen, Cat. No: 12538120). 20 pg of this construct along with 200 ng of tol2 RNA were injected into one-cell stage wildtype embryos and screened for green fluorescence in the vascular endothelial cells. These embryos were raised to adults and in-crossed to generate F1 and F2 generations which were used for our experiments. All embryos were raised at 28.5°C and staged according to the standard guidelines (Kimmel et al., 1995).

To generate the *etv2^Gt(2A-Venus)^* knock-in line, a targeting construct was made by gene synthesis (Genescript Inc) which included *etv2* sgRNA sequence followed by the P2A-*Venus*-pA sequence (Extended Data Fig. 3a) subcloned into the pUC57 vector. To synthesize *etv2* sgRNA, DNA fragment corresponding to *etv2* sgRNA was subcloned into the T7gRNA expression vector (Jao et al., 2013), which was linearized with BamHI and transcribed with T7 RNA polymerase (Promega). *Cas9* mRNA was synthesized from pT3TS-nCas9n vector (Jao et al., 2013) using T3 mMessage mMachine kit (ThermoFisher). A mixture of *Cas9* mRNA, *etv2* sgRNA and the *etv2-2A-Venus* targeting plasmid was injected into wild-type (AB) zebrafish embryos at the 1-cell stage. Embryos showing any specific *Venus* expression were raised through adulthood and screened to identify a founder which produced progeny with a specific expression.

### Whole mount *in situ* hybridization (WISH)

Whole mount *in situ* hybridization was performed as described before(Jowett, 1998). Partial cDNA sequences of *tmem88a* (F: GTACTTTGGAGAAGCTCGCGTC, R: ACCACCAGTAAAGCAACACAGC; Tm **-** 55°C, 421 bp) and *sox7* (F: GTACCTGACGGACACACCTCTC, R: CTCGCAGGATCTCTGAAGACC; Tm **-** 55°C, 479 bp) were amplified from cDNA of 30 hpf embryos using Taq DNA polymerase (M0273, NEB). These amplicons were then sub-cloned into TOPO vector (cat.450640, ThermoFisher) by TA cloning. pExpress-1_egfl7 plasmid was purchased from Dharmacon (clone ID: 7041662). DIG-labeled anti-sense probes were synthesized by digesting corresponding plasmids and transcribed as follows: pSPORT1-etv2(Sumanas et al., 2005), SalI/SP6; pBK-CMV-tal1 (Liao et al., 1998), SalI/T7; TOPO-tmem88a, EcoRV/SP6; TOPO-sox7, EcoRV/SP6 and pExpress1_egfl7, EcoRI/T7. SP6 (cat. no. P1085) and T7 (cat. no. P2075) RNA polymerases were purchased from Promega, Madison, WI.

### RNA fluorescence *in situ* hybridization (RNA-FISH)

*etv2* and *tal1* HCR v3.0 (Hybridization chain reaction) probes were ordered from Molecular Instruments, Inc. Los Angeles, CA. RNA-FISH was performed according to the manufacturer’s protocol and embryos were imaged using a Nikon A1R HD confocal on a TiE inverted microscope with CFI Apo LWD Lambda S 20XC WI NA 0.95.

### Photo-conversion of *fli1a:Kaede* and *etv2:Kaede*

AxioImager compound microscope with Plan-Neofluar 10X/0.3 NA objective (Carl Zeiss Inc., USA) was used for photo-conversion of Kaede. Whole *Tg(fli1a:Kaede)* and *TgBAC(etv2:Kaede)* embryos at 24 hpf were placed in glass depression slides and exposed to DAPI-filtered (405 nm) light for 60 s. Embryos were screened for complete photo-conversion and imaged using a Nikon A1R HD confocal on a FN1 microscope with Nikon CFI75 LWD 16X water dipping objective NA 0.8.

### Photo-activation of *etv2* MO

Photoactivation of *etv2* MO was performed as previously described (Sumanas, 2017). 500 μM of Caging strand (Supernova Life Sciences) and 50 μM of *etv2* MO2 (Sumanas and Lin, 2005) were mixed in 1x Danieau buffer. The mixture was denatured at 70°C for 30 min followed by annealing overnight at 4°C. 2nl of this mixture was injected into zebrafish embryos at 1-cell stage. The mixture and embryos were protected from light by wrapping aluminum foil around the tube or petri dishes. Injections were performed in a dark room with all the light equipped with yellow filters to prevent premature uncaging of *etv2* morpholino. Photoactivation was achieved by exposing injected embryos to 365 nm UV light for 30 min with occasional swirling.

### Identification of SVF cells in Single-cell RNA-Seq analysis (scRNAseq) of zebrafish embryo trunks at 30 hpf

Embryo dissociation and single-cell RNA-seq analysis are described in Metikala *et al* (Metikala et al., 2021) .

To identify SVF cells, the normalized gene expression matrix was filtered based on the following parameters: cells expressing *etv2, tal1* and *lmo2* >1 & *fli1a* <1. Based on these parameters, 23 cells were identified as potential SVF cells.

### Chemical treatments

10 mM metronidazole (MTZ) (M3761, Sigma-Aldrich) solution was made fresh with 0.1% DMSO in embryo medium. Embryos were raised in the presence of MTZ from 50% epiboly until 45 hpf protected from light. A subset of embryos was collected at 45 hpf and fixed in 4% paraformaldehyde (PFA) for WISH analysis. Embryo medium containing MTZ was replaced with fresh embryo medium after washing embryos twice at 45 hpf. Embryos were further raised until 70 hpf for imaging using Nikon A1 confocal microscope or quantified for vascular recovery. Control embryos were raised in the presence of 0.1% DMSO.

Stock solutions were prepared for SU5416 (50 mM) (S8442, Sigma-Aldrich); LDN193189 (10 mM) (SML0559 Sigma-Aldrich) and IWR-1 (10 mM) (I0161, Sigma-Aldrich). For chemical treatment, these stocks were diluted to a final concentration of 10 μM, 20 μM and 20 μM in embryo medium, respectively. Treatment was started at 50% epiboly until 28 or 30 hpf. Embryos were then fixed immediately for WISH.

### Morpholino injections

Morpholinos were obtained from GeneTools LLC (Philomath, OR). *vegfaa* translation blocking (5′- GTATCAAATAAACAACCAAGTTCAT-3′) (Nasevicius et al., 2000), *tal1* splice blocking (5′- AATGCTCTTACCATCGTTGATTTCA-3′) (Dooley et al., 2005) morpholino, *etv2* MO1 (TTGGTACATTTCCATATCTTAAAGT) and *etv2* MO2 (CACTGAGTCCTTATTTCACTATATC)(Sumanas and Lin, 2005) were stored as 3 mM stocks. 10 ng of *vegfaa*, 5 ng of *tal1* and either 5 ng + 5 ng or 10 ng + 10 ng of *etv2* MO1 + MO2 were injected into the yolk at one cell stage and raised until harvesting.

### Imaging and processing

All time-lapse confocal imaging using *TgBAC(etv2:kaede)*, *etv2^Gt(2A-Venus)^*, *etv2^Gt(2A-Venus)^ ; Tg(kdrl:nls-mCherry)*, *etv2^Gt(2A-Venus)^ ; Tg(kdrl:mCherry)* and *etv2^Gt(2A-Gal4)^;UAS:GFP;UAS:mCherry-NTR* embryos was performed using Nikon A1R HD confocal on a FN1 microscope with Nikon CFI75 LWD 16X water dipping objective NA 0.8. Imaging was done at a temperature ranging from 28°C to 30°C. Optical z-slice thickness ranged from 0.2 to 0.5 mm. Images were processed to reduce noise using denoiseAI function and saved as maximum-intensity projections (MaxIP) in NIS-Elements AR software (Nikon Instruments). Movies (.avi) were exported and annotated in Photoshop CC (Adobe).

Confocal imaging using *Tg(fli1a:kaede)* and *etv2^Gt(2A-Gal4)^;UAS:GFP;UAS:mCherry-NTR* embryos was performed using either the same microscope and objective as described above or using Nikon A1R HD confocal on a TiE inverted microscope with CFI Apo LWD Lambda S 20XC WI NA 0.95. Embryos were mounted in 0.6% low melting agarose containing 0.002% Tricaine (Sigma) and 0.003% PTU. Imaging was done at a temperature ranging from 28°C to 30°C. Optical z-slice thickness ranged from 0.2 to 0.5 mm. Images were processed to reduce noise using denoiseAI function and saved as maximum-intensity projections (MaxIP) in NIS-Elements AR software (Nikon Instruments).

etv2^y11^;*Tg(fli1a:GFP)* and *Tg(fli1a:GFP)^y1^* embryos were imaged using an AxioCam MRm monochrome camera (Carl Zeiss) with Zeiss EC Plan-NEOFLUAR 10x/0,3 objective. All WISH embryos were imaged using an AxioCam ICc3 color camera (Carl Zeiss) with Zeiss EC Plan-NEOFLUAR 10x/0,3 objective. All image panels were created using Illustrator CC (Adobe).

### Mouse strains

*Etv2^IRES-H2B-Venus^* mice were a gift from Valerie Kouskoff (Wareing et al., 2012). *Etv2^T2A-iCreERT2/+^* knock-in mice were generated with the help of Biocytogen in C57BL/6J genetic background using CRISPR/Cas9-mediated mutagenesis and 5’- TCGCCTATCAGGATGATATG-3’ gRNA sequence. The sequence encoding T2A-iCreERT2 (Engert et al., 2013; Liu et al., 2017) was inserted using homologous recombination, replacing the TAA termination codon at the 3’end of the *Etv2* gene. The targeting vector encoding T2A-iCreERT2 contained two silent mutations replacing two GAT codons encoding D324 and D325 in the last coding exon of *Etv2* with the synonymous GAC codons. Southern blotting and Sanger sequencing were used to select mice with the correct insertion. Sanger sequencing of the 12-top predicted off-target sites was used to confirm the absence of mutations at these sites. *ROSA^tdTomato/tdTomato^* reporter mice B6.Cg-Gt(ROSA)26Sortm9(CAG-tdTomato)Hze/J were obtained from the Jackson labs (Madisen et al., 2010) (cat# 007909).

#### Tamoxifen injections

To mark *Etv2^+^* cells and their lineage, we mated *Etv2^T2A-iCreERT2/+^* knock-in mice with the reporter *ROSA^tdTomato/tdTomato^* mice. Tamoxifen was dissolved in sesame oil at the concentration of 10 mg/ml and administered by intraperitoneal injection of 100µl of this solution into pregnant females at E9.5. Staining was performed as described below.

#### Mouse embryo whole-mount IF and imaging

Embryos were collected at E10.5, fixed with 4% paraformaldehyde at 4°C overnight, and rinsed in PBS. Embryos were permeabilized in 0.1% Triton (Fisher Scientific, cat# BP151-500) in PBS (PBST) overnight at 4°C and then blocked overnight at 4°C in 10% donkey serum (Sigma, cat# D9663-10ML) in PBST. Embryos were incubated in 1° antibodies in blocking solution for 4 days at 4°C and then washed with PBST. The following 1° antibodies were used: chicken anti-GFP (1:300, Aves, cat# GFP1020), Armenian hamster anti-PECAM1 (1:200, Iowa Developmental Studies Hybridoma Bank Product 2H8 deposited by Bogen, S.A.), mouse anti-ERG (1:1000, Abcam, cat #ab214341), mouse anti-ERG (1:1000, Abcam, cat# ab92513) and rabbit anti-mCherry (1:1000, Abcam, cat # ab167453). Embryos were then incubated with 2° antibodies in blocking solution for 4 days at 4°C. The following 2° antibodies were used at a dilution of 1:300: donkey anti-rabbit 488 (Invitrogen cat# A21206), donkey anti-mouse 555 (Invitrogen cat# A31570), donkey anti-rabbit 555 (Invitrogen cat# 31472), donkey anti-chicken 488 (Jackson ImmunoResearch cat#703-545-155), and goat anti-Armenian hamster 647 (Jackson ImmunoResearch cat # 127-605-160). Nuclei were stained using DAPI (Sigma, cat #32670-5MG-F, 1:1000 dilution of 5 mg/ml stock made in H_2_O). After washes, stained embryos were embedded in the sagittal orientation into 1% agarose (Bio-Rad Laboratories, cat #1613101), dehydrated into 100% methanol and cleared using benzyl alcohol (Sigma, Cat #B-1042)/ benzyl benzoate (Sigma, cat #B-6630), as described(Ramirez and Astrof, 2020). Cleared embryos were placed between two #1.5 coverslips (VWR, cat #16004-312) separated by a rubber spacer (Grace Bio Labs, Cat #664113), as described. Confocal imaging was done using Nikon A1R microscope with 25x CFI Plan Apo Lambda 2 silicone oil objective, N.A. 1.05 (cat # MRD73250). The cardinal vein region of Cre-positive and Cre-negative embryos was imaged posterior to the forelimb.

### Statistical methods

SVF cells were quantified by manually counting the cells on both lateral sides of each embryo. New green-only endothelial cells were quantified manually using ImageJ (v1.52p) Cell Counter plugin. Vascular recovery in MTZ treated embryos was quantified manually in NIS-Elements Viewer (v4.5). Defective vasculature in experiments using photoactivatable etv2 MO was quantified manually in NIS-Elements Viewer (v4.5). Vessels that were more than 75% of normal length were categorized as normal, 25% - 75% as partial defect and less than 25% as severe defect. Calculation of mean ± s.d, two-tailed unpaired t-test, plotting of graphs was performed in GraphPad Prism (v8.0.2).

### Data availability

The original sequence files for zebrafish embryo trunks scRNA-seq at 30 hpf have been deposited to NCBI GEO database under accession number GSE152982 (Metikala et al., 2021).

---

**Figure S1.**
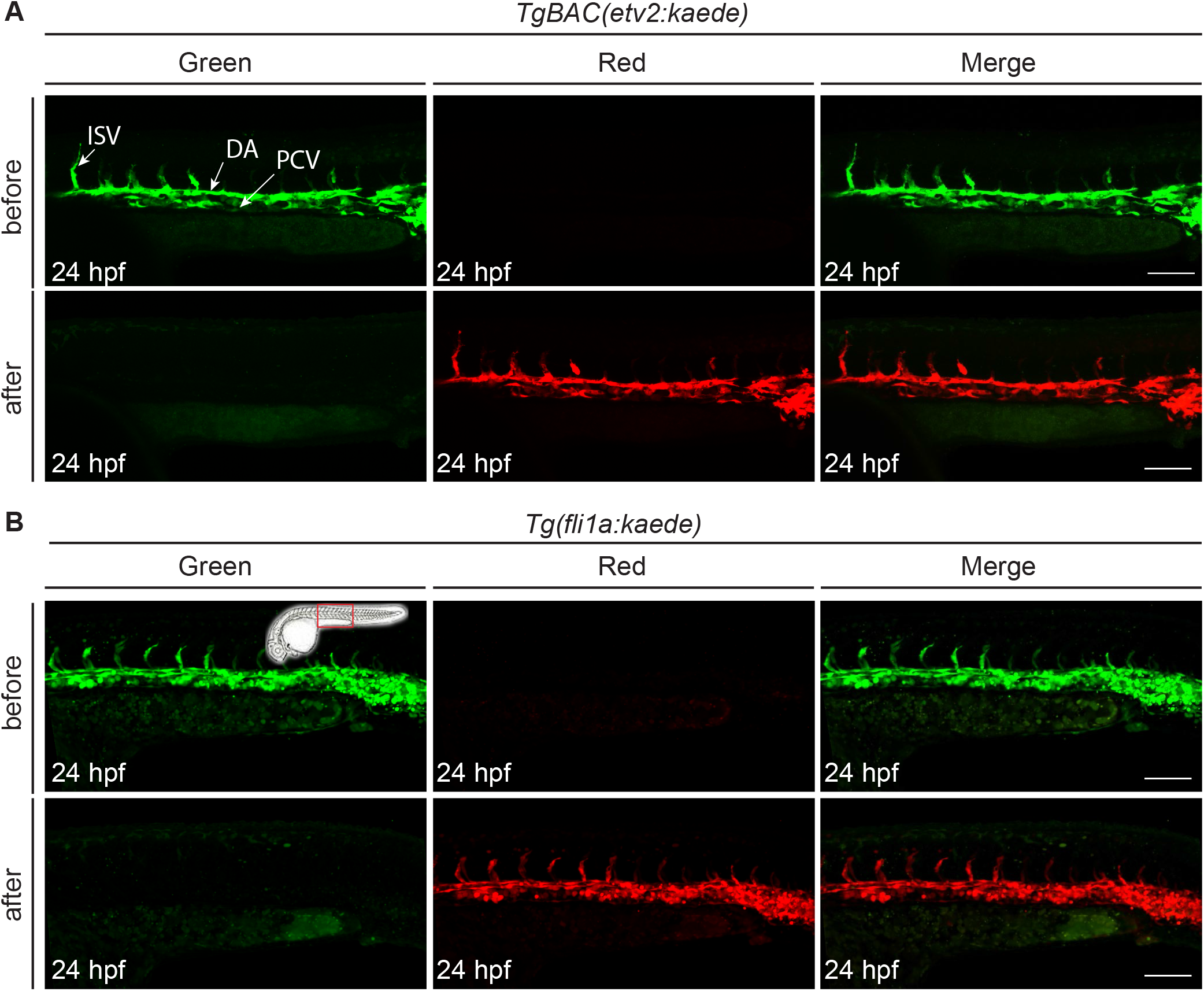
Efficiency of *Kaede* photo-conversion: *Kaede* protein was photo-converted completely from green to red at 24 hpf in (A) *TgBAC(etv2:kaede)* and (B) *Tg(fli1a:kaede)* embryos. The same embryo was imaged in red and green channels before and after photoconversion. PCV, posterior cardinal vein; DA, dorsal aorta; ISV, intersegmental vessels. Scale bars: 100 µm.

**Figure S2.**
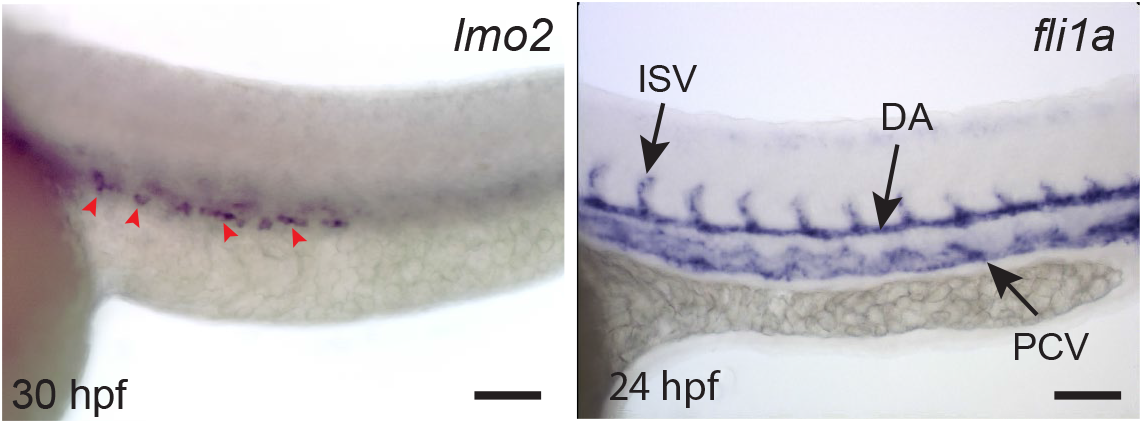
Whole mount *in situ* hybridization analysis of *lmo2* and *fli1a* expression. Note *lmo2* expression in SVF cells (red arrowheads), while *fli1a* is expressed in vascular endothelial cells of the DA, PCV and ISV but not in SVF cells. Lateral view of the trunk region is shown in all images, anterior is to the left. PCV, posterior cardinal vein; DA, dorsal aorta; ISV, intersegmental vessels. Scale bars: 100 µm.

**Figure S3.**
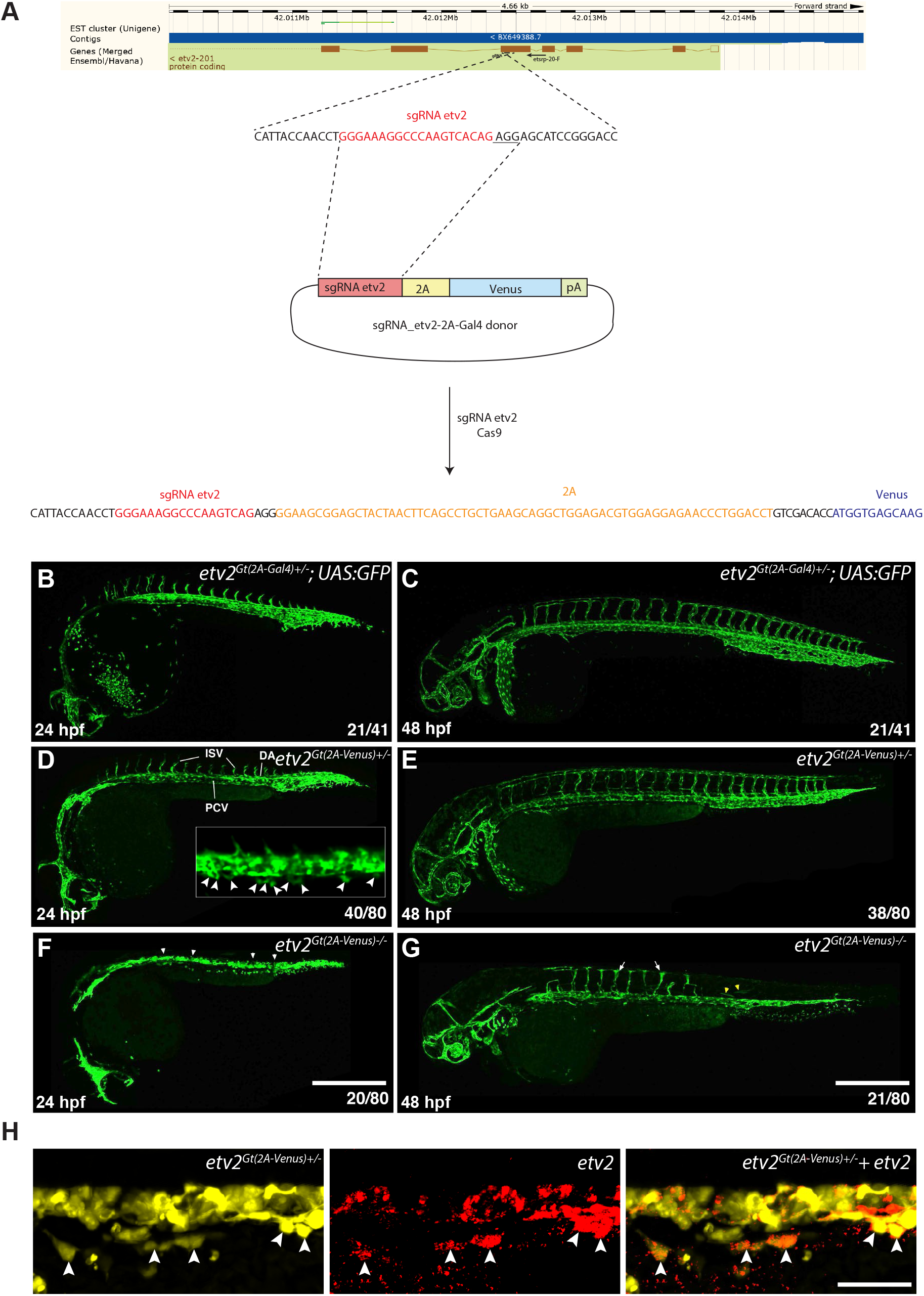
Generation and validation of *etv2^Gt(2A-Venus)^* gene trap line using CRISPR-Cas9 mediated non-homologous recombination. (A) A diagram of *sgRNA-2A-Venus*construct and its insertion into the *etv2*genomic locus. The targeting construct contained *etv2* sgRNA site, followed by in-frame fusion to the viral peptide P2A, the coding sequence of *Venus* and the SV40 polyA sequence. *etv2* sgRNA targets the fifth exon of *etv2* genomic sequence. (B-G) Comparison of *etv2^Gt(2A-Venus)^* and *etv2^Gt(2A-Gal4)^; UAS:GFP* expression at 24 and 48 hpf. Heterozygous embryos in both lines show expression in all blood vessels, including the dorsal aorta (DA), posterior cardinal vein (PCV) and intersegmental vessels (ISV), as well as red blood cells and macrophages. An inset shows a higher magnification view of SVF cells in a *etv2^Gt(2A-Venus)^* embryo (white arrowheads, D). Homozygous *etv2^Gt(2A-Venus)-/-^* embryos show severe defects in vascular development, including the failure of vascular progenitors to coalesce into the axial vasculature at 24 hpf (white arrowheads, F) and absence of intersegmental vessels. Partial ISVs have formed by 48 hpf (arrows, G). *Venus* expression in skeletal muscle cells is also apparent (yellow arrowheads, G). Note that only heterozygous embryos (50%) show fluorescence and appear morphologically normal, while 25% of embryos show mutant phenotype. (H) Fluorescent *in situ* hybridization (FISH) against *etv2* (red) in a *etv2^Gt(2A-Venus)^* embryo (yellow) showing colocalization of *etv2* mRNA and Venus expression in SVF cells (white arrowheads). Scale bars: B: 400 µm, C: 50 µm.

**Figure S4.**
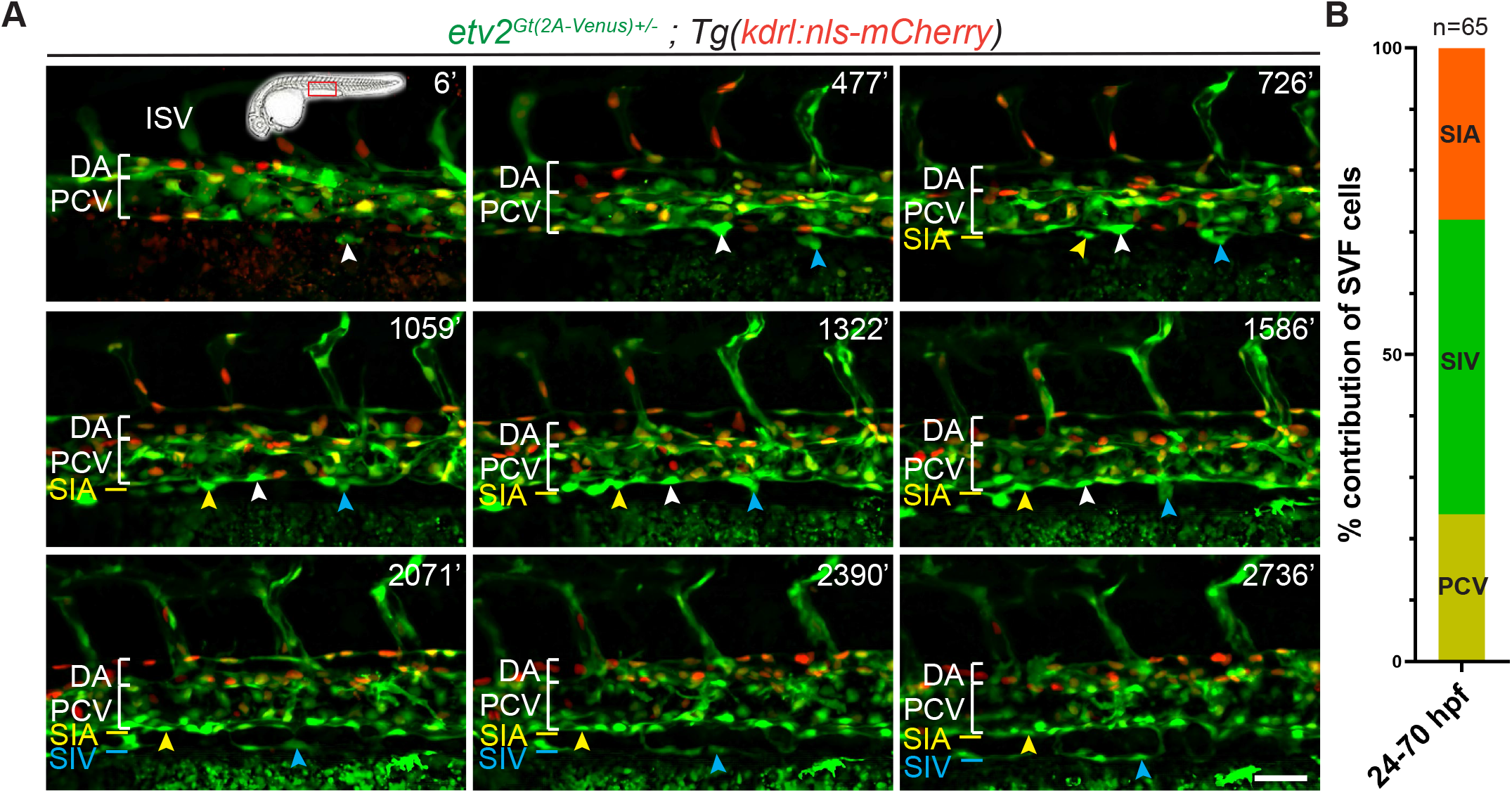
SVF cells contribute to subintestinal vasculature and intercalate into the posterior cardinal vein: (A) Time-lapse images of *etv2^Gt(2A-Venus)+/-^* ; *Tg(kdrl:nls-mCherry)* embryo showing SVF cells which either integrate into the PCV (white arrowheads) or contribute to the formation of SIV (blue arrowheads) and SIA (yellow arrowheads) between 24 and 72 hpf. (B) contribution of SVF cells to various vessels quantified from time-lapse imaging performed at 24-72 hpf (n=6 embryos, 65 SVF cells). Time is shown in minutes after 24 hpf. PCV, posterior cardinal vein; DA, dorsal aorta; ISV, intersegmental vessels; SIV, subintestinal vein; SIA, supraintestinal artery. Scale bars: 50 µm.

**Figure S5.**
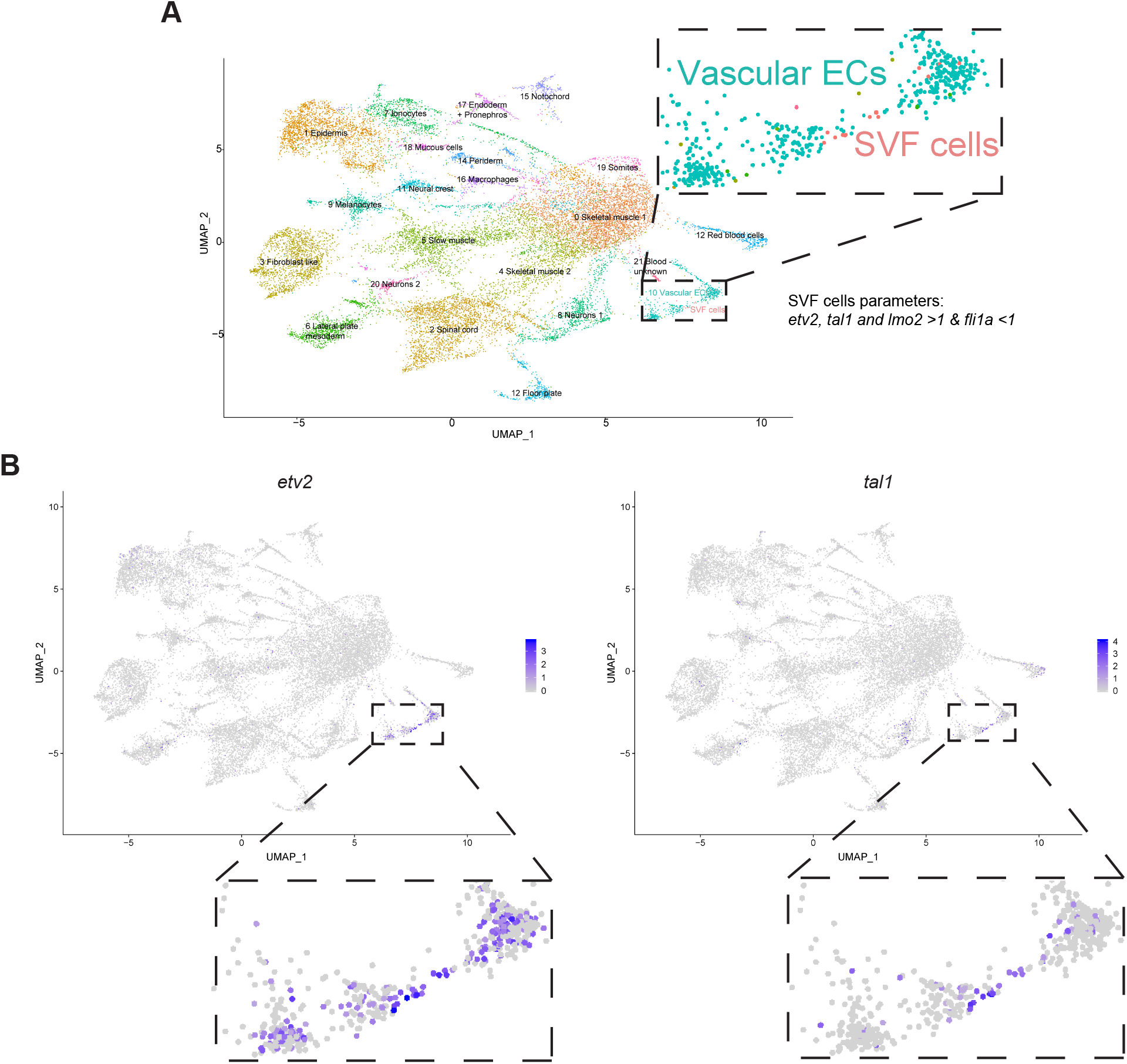
Transcriptome analysis of SVF cells. scRNAseq was performed using cells obtained from zebrafish embryo trunk region at 30 hpf (described in Metikala et al., 2021). (A) Different cell type clusters identified by UMAP analysis. The graph and cluster annotation are based on Metikala et al., 2021. SVF cells were identified based on the differential expression of candidate markers (*etv2, tal1* and *lmo2* >1 and *fli1a*<1). (B) UMAP plots showing relative expression of *etv2* and *tal1* (log2 scale) in all the clusters and in vascular endothelial cells (insets). Note that SVF cells have the highest expression of *etv2* and *tal1*.

**Figure S6.**
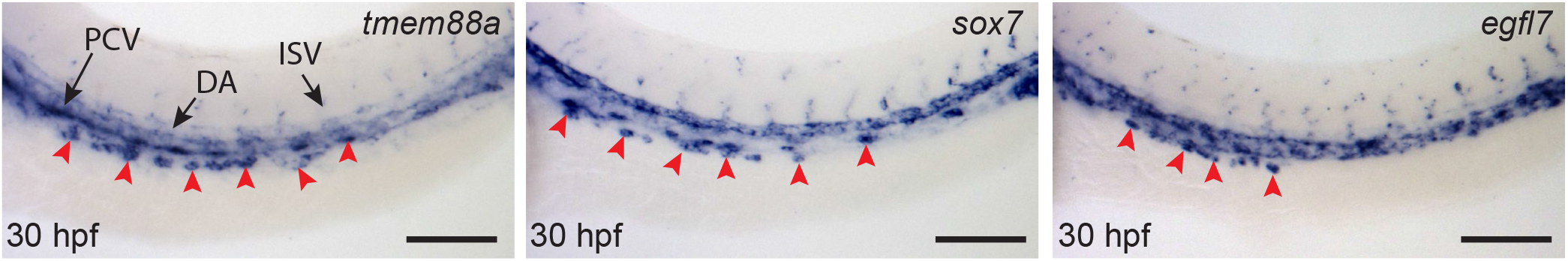
Validation of SVF marker genes: WISH analysis of *tmem88a*, *sox7* and *egfl7* expression in wt embryos at 30 hpf. Note expression in SVF cells (red arrowheads) and blood vessels. Dorsolateral view of the trunk region is shown in all images, anterior is to the left. PCV, posterior cardinal vein; DA, dorsal aorta; ISV, intersegmental vessels. Scale bars: 100 µm.

**Figure S7.**
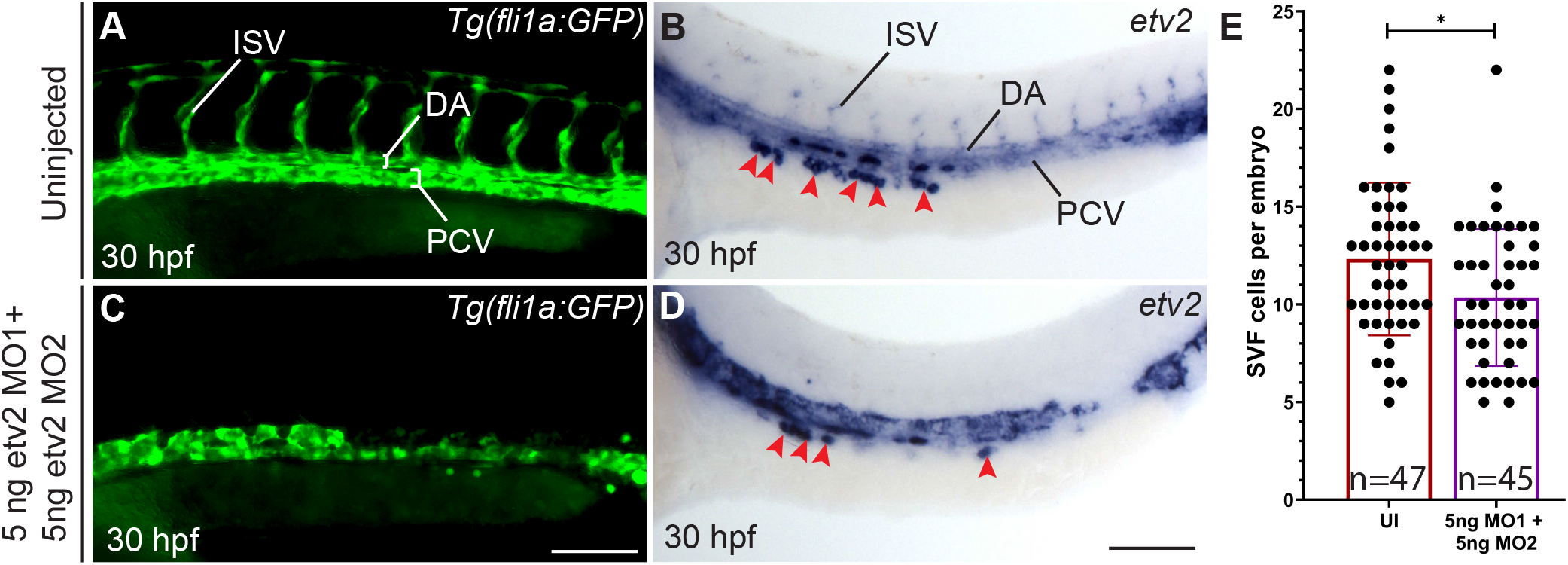
*etv2* knockdown has only minor effect on the specification of SVF cells: *Tg(fli1a:GFP)* embryos were injected with 5 ng of *etv2* MO1 + *etv2* MO2 at one-cell stage. (A,C) *etv2* morphants show defective axial vasculature formation and absence of ISVs at 30 hpf. (B,D) WISH for *etv2* expression shows defective axial vasculature. Note that SVF cells (red arrowheads) are still present although their number is slightly reduced in *etv2* knockdown embryos (12.32 SVF cells per embryo) compared to controls (10.36 SVF cells per embryo). Due to defects in the formation of axial vasculature it is challenging to distinguish SVF cells from vascular progenitors of axial vessels. (E) Quantification of SVF cells in *etv2*morphants and control uninjected embryos at 30 hpf. mean ± s.d, *P<0.05 (two-tailed unpaired t-test). Each data point represents one embryo. Data from 2 independent experiments. PCV, posterior cardinal vein; DA, dorsal aorta; ISV, intersegmental vessels; UI, uninjected embryo. Scale bars: 100 µm.

**Figure S8.**
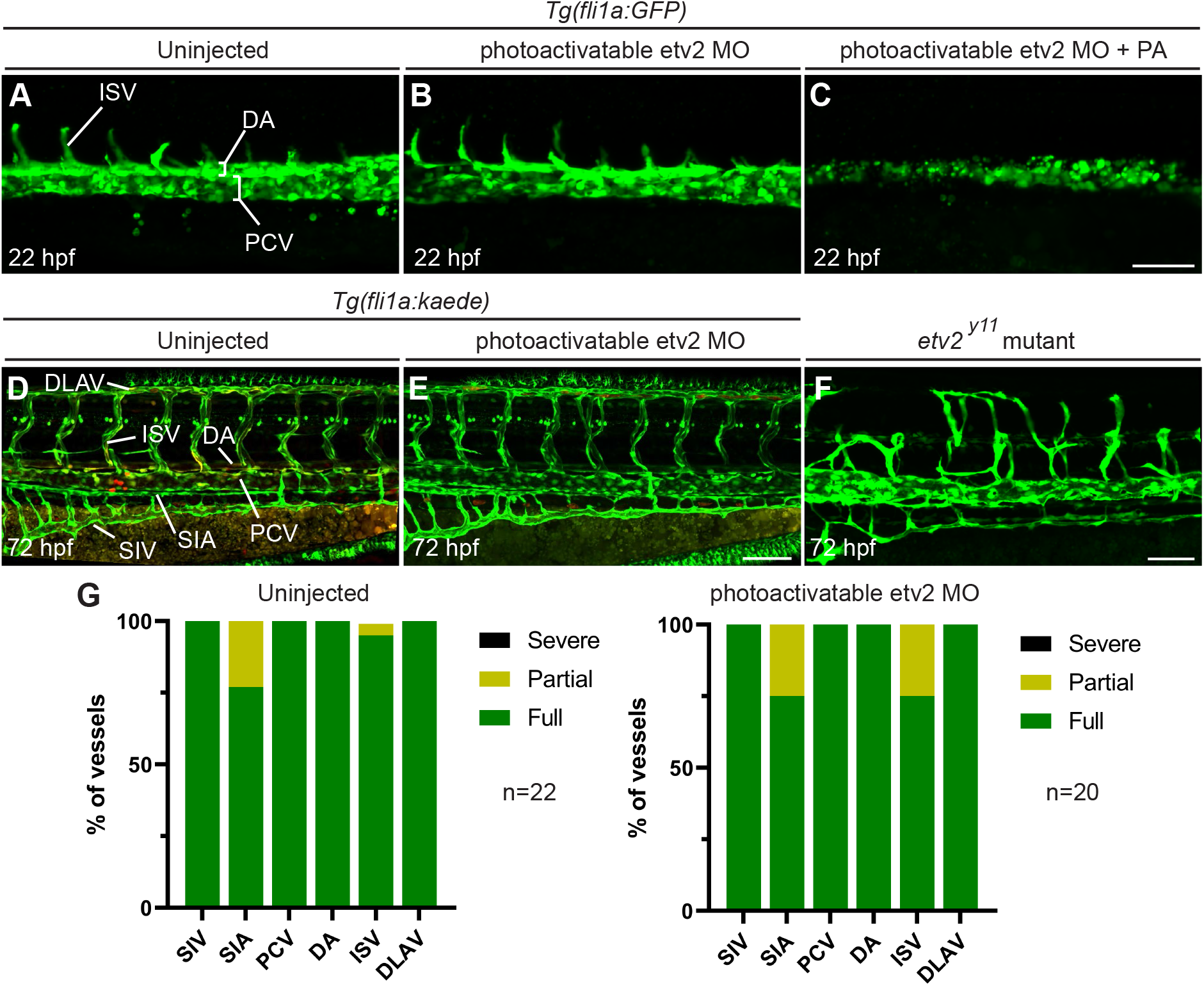
Validation of photoactivatable *etv2* MO: (A-C) Embryos were injected with photoactivatable *etv2* MO in *Tg(fli1a:GFP)* embryos at one-cell stage followed by *etv2* MO photoactivation (PA) at 50% epiboly (6 hpf). Defective axial vasculature formation was apparent in injected and photoactivated embryos at 22 hpf (C) but not in embryos that were injected but never photoactivated (B). (D,E) *Tg(fli1a:Kaede)* embryos were injected with photoactivatable *etv2* MO at one-cell stage but never photoactivated. Morphants showed normal axial vessel formation without any defects in the SIA formation at 72 hpf. (F) *etv2^y11^* mutant embryo showing strong defects in vessel formation at 72 hpf. Note the absence of SIA formation. (G) quantification of fully formed, partially affected, or severely affected vessels in the trunk in uninjected control embryos (n=22) and *etv2* morphants (n=20). Data from 2 independent experiments. PCV, posterior cardinal vein; DA, dorsal aorta; ISV, intersegmental vessels; SIA, supraintestinal artery DLAV, dorsal longitudinal anastomosing vessel; SIV, subintestinal vein. Scale bars: 100 µm.

**Figure S9.**
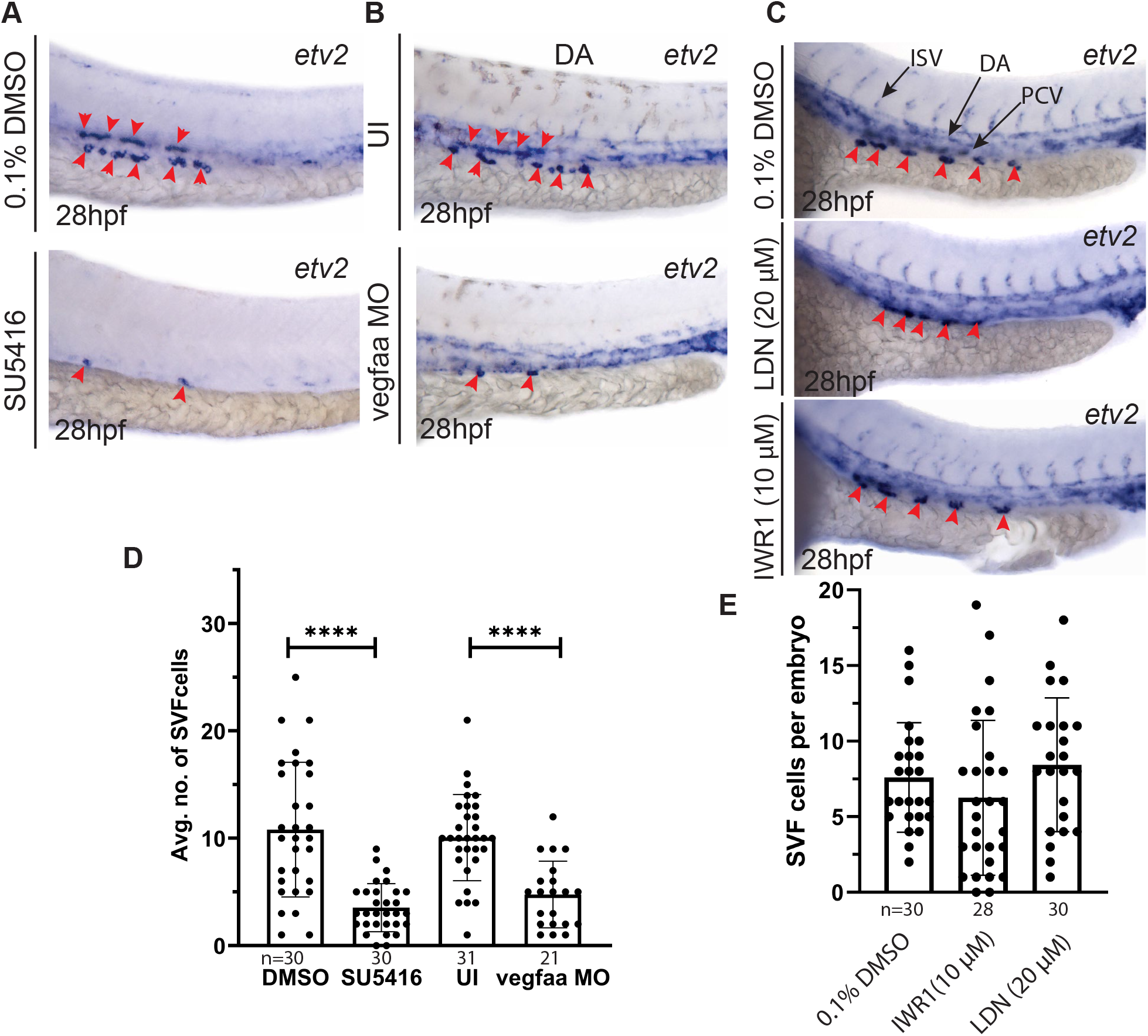
*Vegf* signaling pathway, but not *wnt* and *bmp* pathways, is necessary for the formation of SVF cells: (A) WISH analysis for *etv2* expression at 28 hpf in SU5416 treated embryos and DMSO controls, (B) WISH analysis for *etv2* expression at 28 hpf in *vegfaa* MO injected embryos and uninjected controls. (C) Embryos were raised in the presence of *wnt* inhibitor, IWR1, and *bmp* inhibitor, LDN193189, from 50% epiboly to 28 hpf. WISH was performed for *etv2* showing expression in vasculature and SVF cells (red arrowheads) (D) Quantification of SVF cells (red arrowheads) in control DMSO treated (n=30) and SU5416 treated (n=30) embryos, control uninjected (n=31) and *vegfaa* morpholino injected embryos (n=21). (E) Quantification of SVF cells (red arrowheads) in wildtype siblings (n=53) and *npas4l^-/-^* embryos (n=61). Data from 3 independent experiments. mean ± s.d, \*\*\*\**P*<0.0001 (two-tailed unpaired t-test). Each data point represents one embryo. PCV, posterior cardinal vein; DA, dorsal aorta; ISV, intersegmental vessels. Scale bars: 100 µm.

**Figure S10.**
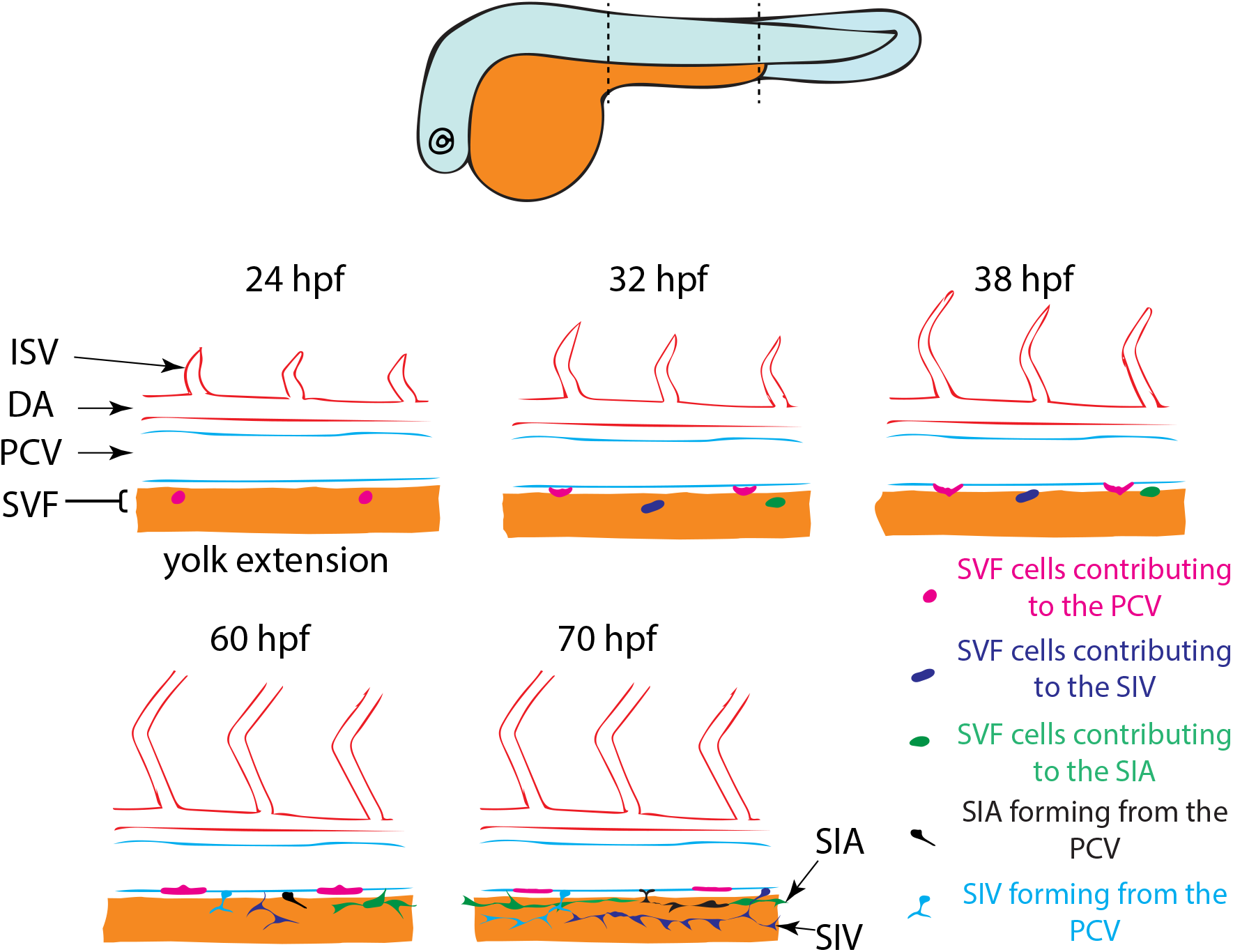
SVF cell contribution to the axial vessels: The illustration shows SVF cells contribution to the PCV, SIV and SIA. Some of these cells contribute to the PCV while others form a part of the SIV and SIA. Note that all three vessels have partial contributions from SVF cells, and the rest of the SIA and SIV are derived from non-SVF cells located in the PCV as it has been previously demonstrated (Hen et al., 2015 and Koenig et al., 2016).

**Table S1:** Lineage tracing of SVF cell contribution using *etv2^Gt(2A-Venus)+/-^* crossed to *Tg(kdrl:nls-mCherry)* or *Tg(kdrl:mCherry)* reporter lines. Time-lapse imaging was performed from 24 to approximately 70 hpf. n = 6 embryos

| SVF cell S.No. | Final contribution | No. of cells after proliferation |
| --- | --- | --- |
| 1 | PCV | 2 |
| 2 | PCV | 1 |
| 3 | PCV | 2 |
| 4 | PCV | 2 |
| 5 | PCV | 2 |
| 6 | PCV | 3 |
| 7 | PCV | 2 |
| 8 | PCV | 2 |
| 9 | SIV | 4 |
| 10 | SIV | 2 |
| 11 | SIV | 4 |
| 12 | SIV | 2 |
| 13 | SIV | 1 |
| 14 | SIV | 2 |
| 15 | SIV | 7 |
| 16 | SIV | 2 |
| 17 | SIV | 3 |
| 18 | SIV | 4 |
| 19 | SIA | 2 |
| 20 | SIA | 1 |
| 21 | SIA | 1 |
| 22 | SIA | 4 |
| 23 | SIA | 1 |
| 24 | SIA | 1 |
| 25 | SIA | 5 |
| 26 | SIA | 3 |
| Total no. of cells |  | 65 |

**Table S2:**
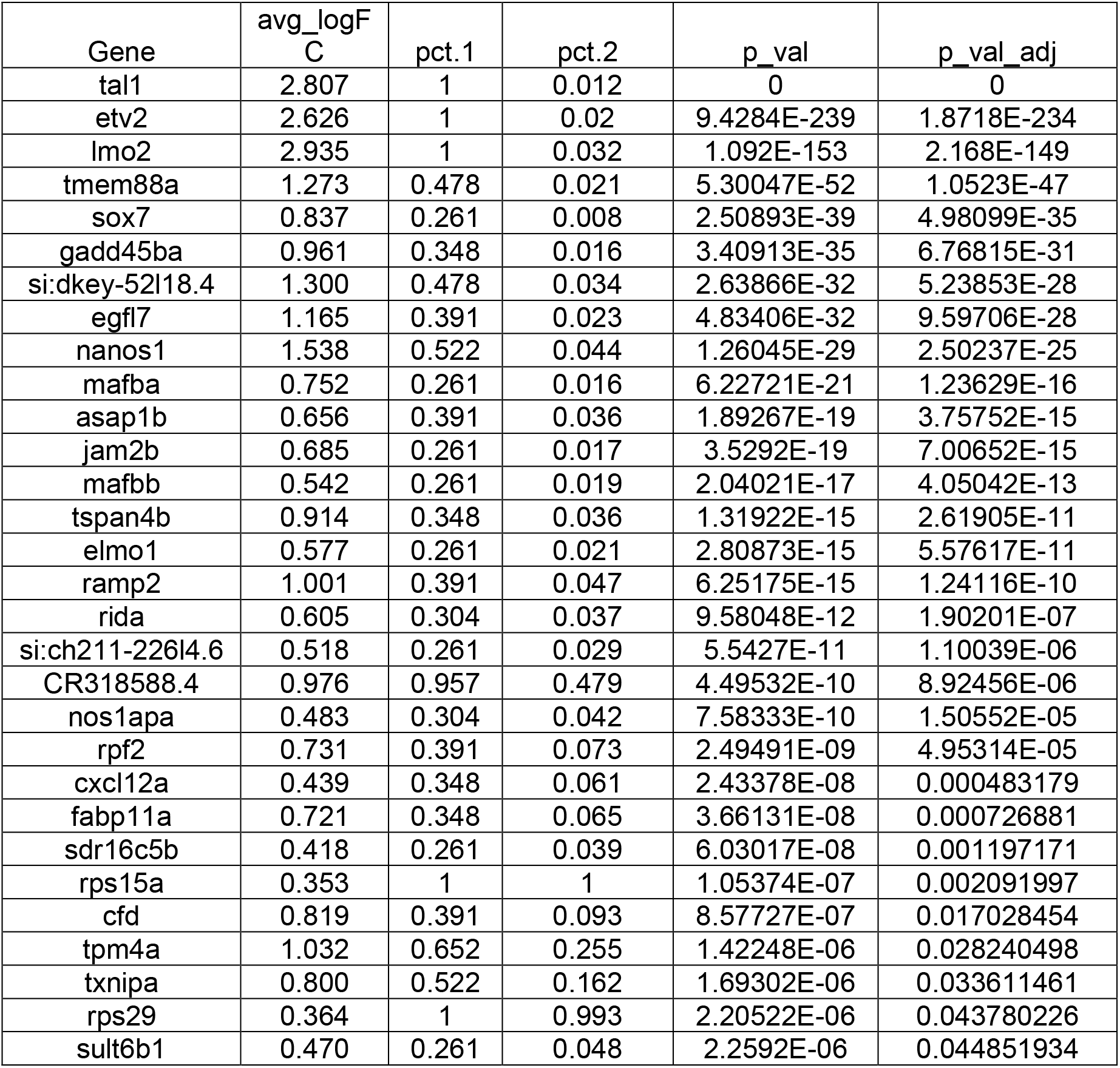
List of top genes expressed in SVF cells obtained using zebrafish embryo trunks at 30 hpf.

**Table S3:**
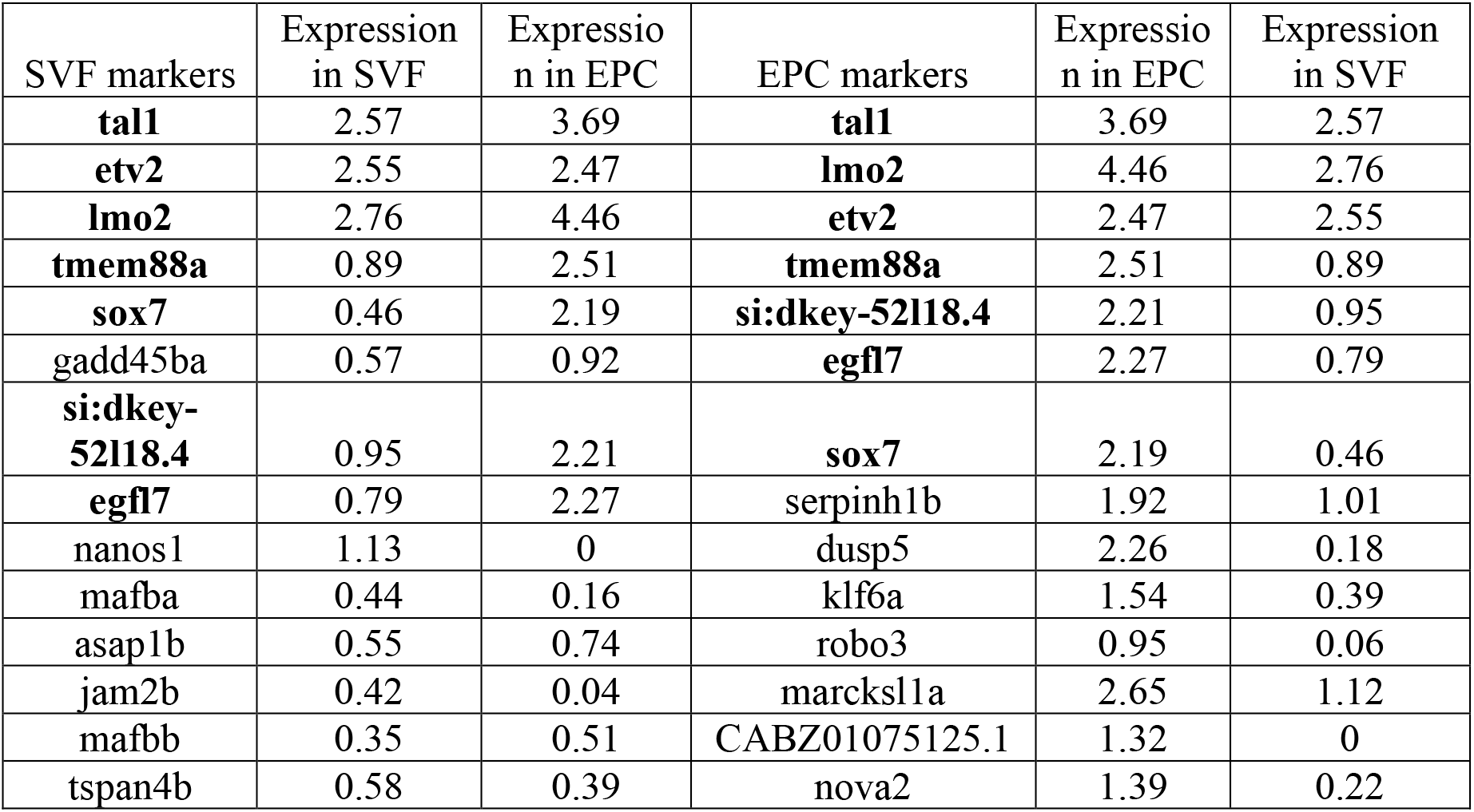
Comparison of relative top marker expression in SVF cells (30 hpf, trunk scRNA-seq) and EPC cells (20-somite stage, *etv2^Gt(2A-Gal4)^* line scRNA-seq). Log2 differential expression values are shown (compared to all other cell types). Shared top marker genes are marked in bold. EPC data from Chestnut et al 2020.

## Supplemental Video Legends

**Video S1. 3D volume reconstruction of SVF cells and pronephros:** *Tg(enpep:GFP)* embryo which labels pronephros was stained by FISH against *etv2* (red) and imaged using confocal microscopy at 30 hpf. SVF cells (red) are located either medially, laterally or ventrally adjacent to the pronephros (green). The video initially shows external dorso-lateral view and rotates to show the internal view. The video ends by showing the z-axis build down (going through each z-section from the external to the internal view) (see Figure 2F,G).

**Video S2. Integration of SVF cells into the PCV:** Time-lapse video of *etv2^Gt(2A-Venus)+/-^* ; *Tg(kdrl:mCherry)* embryo showing multiple SVF cells integrating into the PCV (white arrowheads). Embryo was imaged from 24 hpf to 40 hpf. PCV, posterior cardinal vein; DA, dorsal aorta; Scale bar: 100 µm.

**Video S3. Integration of SVF cells into the PCV:** Time-lapse video of *etv2^Gt(2A-Venus)+/-^* embryo showing integration of SVF cells into the PCV. Two SVF cells originated outside of vasculature, migrated towards the PCV and intercalated into it (white arrowheads). Embryo was imaged from 24 to 44 hpf. PCV, posterior cardinal vein; DA, dorsal aorta. Scale bars: 50 µm.

**Video S4. Mechanism of integration of an SVF cell into the PCV:** Time-lapse video of *etv2^Gt(2A-Venus)+/-^* ; *Tg(kdrl:mCherry)* embryo showing the events in the integration of SVF cell into the PCV. Two SVF cells can be seen migrating towards the PCV and making contact with the ECs in PCV. This is followed by the rearrangement of ECs in the PCV and integration of SVF cells into the PCV. Subsequently these SVF cells initiate *kdrl:mCherry* expression and undergo proliferation (white arrowheads) (see Figure 3B). Embryo was imaged from 24 hpf. PCV, posterior cardinal vein; DA, dorsal aorta; ISV, intersegmental vessel. Scale bar: 50 µm.

**Video S5. Contribution of SVF cells to the trunk vasculature after 24 hpf:** Time-lapse video of *etv2^Gt(2A-Venus)+/-^* ; *Tg(kdrl:nls-mCherry)* embryo showing the emergence of SVF cells, their migration and incorporation into the PCV (white arrowheads), or contribution to the SIV (blue arrowheads) and SIA (yellow arrowheads) (see Figure S4A). Embryo was imaged from 24 to 70 hpf. PCV, posterior cardinal vein; DA, dorsal aorta; ISV, intersegmental vessels; SIV, subintestinal vein; SIA, supraintestinal artery. Scale bars: 50 µm.

**Video S6. Vascular recovery after endothelial cell ablation:** a-c, time-lapse videos of *etv2^Gt(2A-Gal4)+/-^;UAS:GFP;UAS:mCherry-NTR* embryo showing vascular recovery. All *etv2* expressing endothelial cells were ablated with MTZ treatment at 6-45 hpf and the vasculature was let to recover. Initial location of cells expressing only GFP and final location of cells co-expressing both GFP and mCherry are marked in dotted lines in both channels and merge. Recovered vessels were observed in putative PCV and SIV locations. It is important to note that recovered VECs initially expressed GFP followed by mCherry which has a longer maturation period. Embryo was imaged from to 47 to70 hpf. PCV, posterior cardinal vein; DA, dorsal aorta; SIV, subintestinal vein. Scale bar: 50 µm.

